# Influenza H1N1 Imprinting Uncouples Serological and B-Cell Responses in Mice

**DOI:** 10.64898/2026.10.26.761834

**Authors:** Elias Tjärnhage, Khang Lê Quý, Taissa de Matos Kasahara, Marit Lovise Toven Dæhli, Victor Greiff, Gunnveig Grødeland

## Abstract

Antigenic drift reduces the protective potential of antibodies generated against earlier influenza variants. Unfortunately, a new infection or vaccination often preferentially recalls antibodies targeting previously encountered strains, thereby limiting responses to newly emerged variants. Although this phenomenon, known as immunological imprinting, has been well documented at the serological level, its underlying cellular mechanisms remain poorly understood. To resolve imprinting at both the serological and cellular level, we combined serum antibody profiling with targeted B-cell characterization in mice, including single-cell B-cell receptor (BCR) sequencing of germinal center (GC) B cells following secondary H1N1 challenge and bulk BCR sequencing of bone marrow plasma cells. We found that a primary infection with the 2009 pandemic H1N1 (“swine flu”) strain largely prevented the generation of new serum antibodies following secondary exposure to the antigenically distinct H1N1 strain A/Puerto Rico/8/1934 (PR8), whereas the reciprocal infection sequence updated antibody specificities towards the last encountered virus. GC B cells did not mirror serological imprinting and instead generated updated BCR repertoires directed against the heterologous challenge strain. Likewise, BCR diversity in draining lymph node B cells failed to mirror the serological imprinting observed in the antibody response. Together, these findings reveal a striking uncoupling of cellular and serological immune responses during influenza imprinting, highlighting the need to study both compartments to better understand immune memory and guide the development of improved seasonal influenza vaccines across a diverse population.

## Introduction

Seasonal influenza viruses keep challenging our immune system with new variants yearly, and the neutralizing antibodies generated against last year’s strain may no longer be effective against this year’s antigenically drifted variant. In the 1950’s, it was observed that the first influenza strain encountered by an individual would dominate later responses^1^. High serum levels of antibodies specific for the first strain would be maintained even after repeated exposures to antigenically drifted strains. A few years later, Dr. Thomas Francis named this concept “Original Antigenic Sin”^2^, and we now know that such imprinting will influence an individual’s susceptibility to seasonal influenza infections^3^. Imprinting have been observed for many different circulating influenza subtypes^4–7^, and where responses against the surface protein HA are more affected^8^. That said, most of our current understanding on the imprinting effect from the first influenza exposure is based solely on serological evaluations^9,10^.

High affinity antibodies represent one of the products emanating from the GC reaction. GC B cells are clonally selected for affinity to a relevant antigen, and may undergo several rounds of somatic hypermutation (SHM) before surviving clones exit as long-lived antibody secreting cells or memory B cells (MBC)^11^. For viruses with a variable antigenic profile, such as influenza, we still do not fully understand whether already established MBC would re-enter the GC and further mutate their BCR upon secondary exposure to an antigenically drifted variant, or if they would rapidly transition into antibody secreting cells with specificity towards the imprinted strain. We know that certain MBC markers and class-switching can influence this fate^12,13^, and that the similarity between the strain first encountered and the subsequent strain affects the specificity of the produced antibodies^14^. The more different the new virus is from previous exposures, the more likely it is for the initial imprinting to be overcome^15^.

Here, we set out to examine the link between antibody imprinting and the underlying B cell response in a system where influenza exposures could be controlled. Thus, we coupled serum antibody responses in mice first exposed to either influenza A/California/07/2009 (H1N1) (CA07) or PR8 viruses to B cell evaluations and V(D)J sequencing of single HA specific B cells from draining LN, and large-scale BCR sequencing of bone marrow PCs. Interestingly, while primary CA07 infection prevented the formation of PR8-specific antibodies following re-infection with PR8, the inverse was not true. At the cellular level, we observed a more balanced induction of de novo B cell specificities following heterologous re-infection, and with no clear convergence towards unique CDR sequences for either the HA(CA07) or the HA(PR8) specific B cells. In sum, we found that the serological phenomenon of immune imprinting will differ between influenza strains, and that it could not be explained by similar restraints at the B cell level.

## Results

### CA07 priming suppresses formation of antibodies against PR8 upon secondary infection, whereas PR8 does not exhibit similar suppressive potential

To evaluate the imprinting effect of primary influenza infection on secondary responses, BALB/c mice were sequentially challenged with sublethal doses of PR8 and CA07 at a 6-week interval. Please note that HA of PR8 and CA07 have an overall amino acid sequence identity of ∼82%, while the globular head that is the prime site for binding of neutralizing antibodies, the HA1 subunits, have a sequence identity of ∼76% (Fig. 1A). Blood samples were collected longitudinally, and draining LN and bone marrow collected two weeks post the secondary exposure (Fig. 1B).

**Figure 1.**
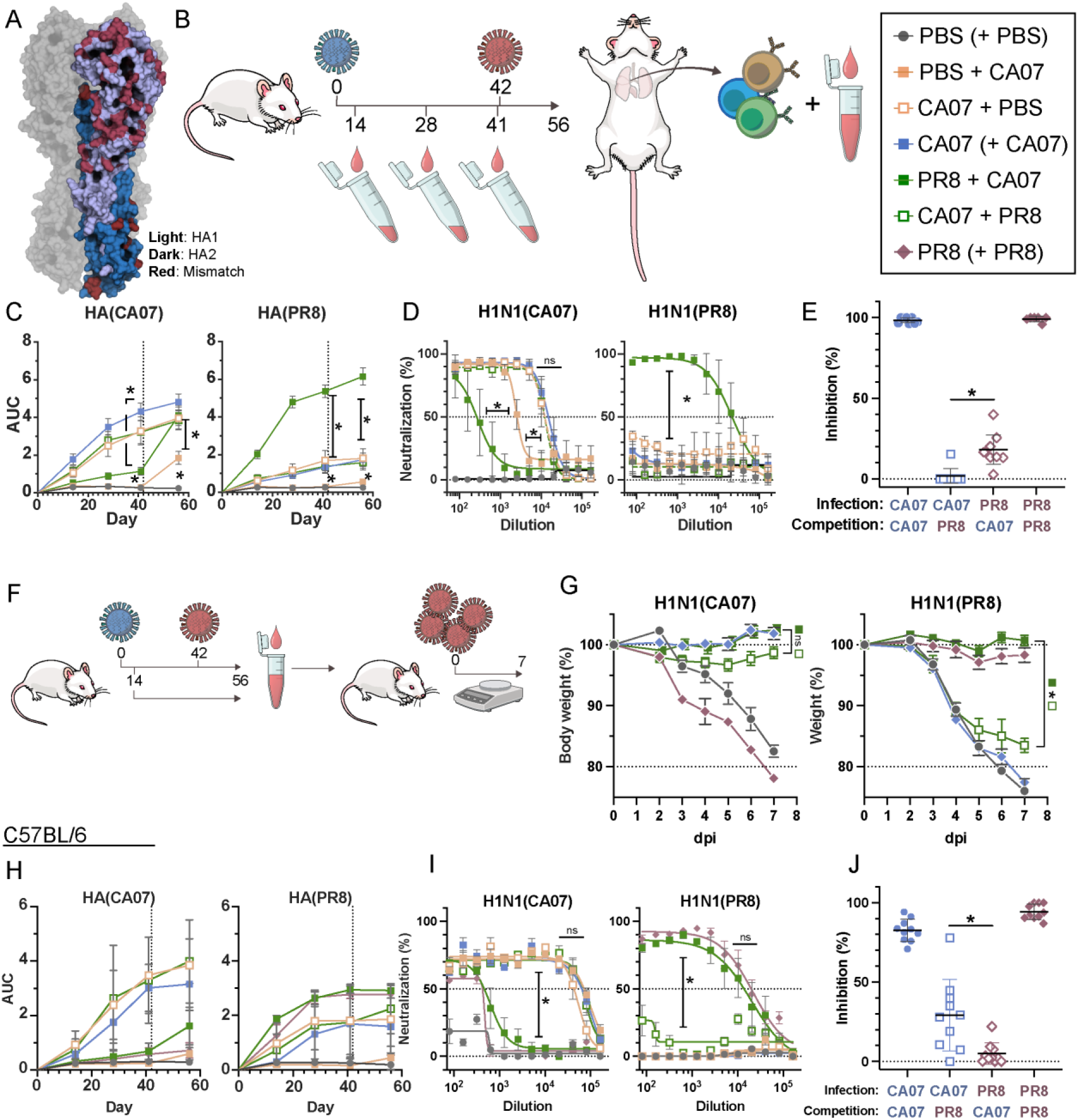
Priming with CA07 will impair formation of new responses against PR8 in BALB/c and C57BL/6 mice. **A)** Structural representation of influenza HA monomer from A/Puerto Rico/8/1934(H1N1) (PR8) and A/California/07/2009(H1N1) (CA07), with amino acid differences highlighted in red. The head domain (HA1) is depicted in light, and the stem domain (HA2) in dark. There is a ∼76% identity between HA1 domains and ∼82% between complete HA from PR8 and CA07 (PDB ID PR8: 1RUZ, CA07: 3AL4). **B)** Experimental setup. In brief, BALB/c mice were infected with a sublethal dose of either PR8 or CA07 at day 0, followed by a secondary challenge at day 42. Challenge order and strains are depicted for each experimental group (right). Blood samples were collected at the indicated days post infection (dpi). At day 56, mediastinal LNs and spleens were collected for B cell evaluation. **C)** Sera were evaluated for IgG binding to HA(CA07) (left) and HA(PR8) (right) in ELISA. Stippled line denotes timepoint for re-infection. Data was pooled from five separate experiments (Extended data Fig. 1, Experiments A-E), n=34 for PBS+PBS and CA07+CA07; n=28 for PBS+CA07 and PR8/CA07, and n=12 for CA07+PBS and CA07+PR8. Data is depicted as Mean±95%CI. **D)** Microneutralization against H1N1(CA07) (left) and H1N1(PR8) (right) with sera collected at day 56. Stippled line indicates cut off for positive neutralization activity. Data was pooled from three separate experiments (Extended data Fig. 1, Experiments A-C): n= 18 for PBS/PBS and CA07/CA07; n=12 for PBS/CA07, CA07/PBS, and PR8/CA07. Data is depicted as Mean±95%CI. **E)** Competition ELISA with serum collected 41 days after a single infection with either CA07 or PR8 (n=8 per infection) (Extended data Fig. 1, Experiment E). Sera were pre-adsorbed with competitor HA before being added to microtiter plate. Data is depicted as normalized signal loss, Mean±95%CI. **F)** Illustrative overview of experiment with serum transfer. In brief, sera collected post two sublethal influenza infections at day 56 were pooled within groups, and administered i.v. to naïve mice. The next day, these mice were infected with a lethal dose (2×LD_50_)of either influenza CA07 or PR8. **G)** Weight loss after serum transfer and lethal influenza infections, n=8 mice per group. Data depicted as Mean±SEM, and statistical significance indicated for day 7. **H-J)** Experiment outlined above in B was repeated for C67BL/6 mice. **H)** Serum IgG against HA(CA07) (left) and HA(PR8) (right) (n=5 mice/group). **I)** Serum neutralization against CA07 (left) and PR8 (right) (n=5 mice/group). **J)** Competition ELISA with sera collected at day 41 post a single infection (n=10 mice/group). Statistical significance was calculated by Kruskall-Wallis’s one-way ANOVA, (C-D, G-I) and Mann-Whitney test (E, J), with a False Discovery Rate at Q=0.05.

An examination of serum antibody responses showed that a primary challenge with either CA07 or PR8 could significantly raise high homologous serum IgG levels, with some lower levels of cross-reactive responses observed against the other strain (p<0.0025). Interestingly, while priming with CA07 followed by a PR8 re-infection did not elevate antibody responses against PR8 (p=0.8864), a significant increase (p<0.001) was observed in mice primed with PR8 followed by CA07 re-infection (Fig. 1C, and Extended data Fig. 1). The same trend was observed when restricting the evaluation of serum reactivity to the HA1 subunit of HA (Extended data Fig. 1M), and there were no significant IgG1 and IgG2a subclass differences observed for the differently primed groups (Extended data Fig. 1B, E, H). The neutralizing capacity of the collected sera corresponded well with the observed IgG levels in that only serum from PR8 imprinted mice were able to neutralize PR8 virus (p<0.001). Mice primed with CA07 and re-infected with PR8 failed to do so. By contrast, mice primed with PR8 and then re-infected with CA07 could neutralize CA07 virus, albeit at a lower level than one or two exposures to CA07 (p<0.001) (Fig. 1D, and Extended data Fig. 1C, F, I). Importantly, we observed significantly higher levels of cross-reactive antibodies in mice imprinted with PR8 as compared to CA07 by pre-adsorbing sera from the different groups with soluble HA before detecting remaining IgG in a competition ELISA (p<0.001) (Fig. 1E). As a result, we subsequently tested whether the in vivo protective capacities of the induced antibodies were replicating observations from the ELISA and microneutralization assays, and performed an experiment where sera from the different groups were adoptively transferred to naïve mice prior to lethal influenza challenges with either CA07 or PR8 (Fig. 1F). In correspondence with above data, results demonstrated that both sera from mice sublethally infected with first CA07 and then PR8, or vice versa, protected against a lethal dose of CA07, but mice first exposed to CA07 followed by PR8 were not protected against PR8 (Fig. 1G). As expected, sera from repeat infections with either PR8 or CA07 could protect against the corresponding homologous strain.

Could it be that the dominance observed for CA07 priming was unique for BALB/c mice? To test this, we evaluated sequential infections also in C57BL/6 mice that genetically harbor a different baseline immune repertoire^16^. Results demonstrated also here that priming with CA07 suppressed antibody responses to PR8 upon secondary infection, whereas priming with PR8 allowed formation of antibodies against CA07 upon secondary exposure (Fig. 1H, I). However, we observed a difference between BALB/c and C57BL/6 when assessing cross-reactivity after primary infections with either CA07 or PR8 in competition ELISA. In C57BL7/6 mice, CA07 raised significantly more cross-reactive IgG than PR8 (p=0.004) (Fig. 1J), whereas the opposite was true for BALB/c (Fig. 1E). It is likely that this was caused by either differences in BCR repertoire ^17,18^, MHC haplotype, or a combination of both.

Different influenza viruses have different potentials for infecting mice. Whereas PR8 is a mouse adapted strain that has been used for decades in research, CA07 is a human influenza strain that can also raise infection in mice (but a larger dose is typically needed, as compared to PR8). To assess if the observed CA07 dominance over a subsequent PR8 infection was due to dose or other virus properties, we set up an experiment where mice were either primed as before with CA07 or vaccinated with HA from CA07. Of note, the vaccine format used was DNA, and where HA was genetically linked to a gene encoding a scFv specific for MHCII molecules, ^19,20^. We know from before that such MHCII specific targeting of antigen will equalize the induced levels of IgG responses for different influenza subtypes and strains ^21,22^. In brief, both the primary vaccination and infection resulted in robust antibody responses and formation of MBC against CA07 (Extended data Fig. 2 A-B). Importantly, results further demonstrated that priming with either the sublethal CA07 infection or vaccination similarly suppressed formation of antibodies against PR8 upon secondary exposure (Extended data Fig. 2, C-F). As such, intrinsic properties of CA07 HA must have caused the downstream suppression of PR8 specific antibodies after secondary infection, and not viral differences in dose or immunogenicity.

### Serological imprinting is not replicated in GC B cells

To assess the cellular foundation for the observed serological imprinting, we evaluated GC B cells following sublethal viral infections with PR8 and CA07 (Fig. 2A, Extended data Fig. 3). In contrast to the cross-reactive antibodies observed after priming with either PR8 or CA07 (Fig. 1E, J), we observed no corresponding cross-reactive GC B-cells at day 28 post primary viral challenge in BALB/c mice (Fig. 2B, Extended data Fig. 4, A-B). Upon secondary infections with heterologous strains, we detected solid populations of GC B-cells against both the priming strain and the strain used for secondary exposure (Fig. 2C, and Extended data Fig. 4, C-E). Only limited numbers of cross-reactive GC B-cells were detected, although with an increase after two heterologous infections as compared to two homologous infections with either PR8 or CA07. Interestingly, there was a higher percentage of PR8-specific GC B-cells observed in mice that had been primed with CA07 and re-infected with PR8, as compared to the opposite (Fig. 2D).

**Figure 2.**
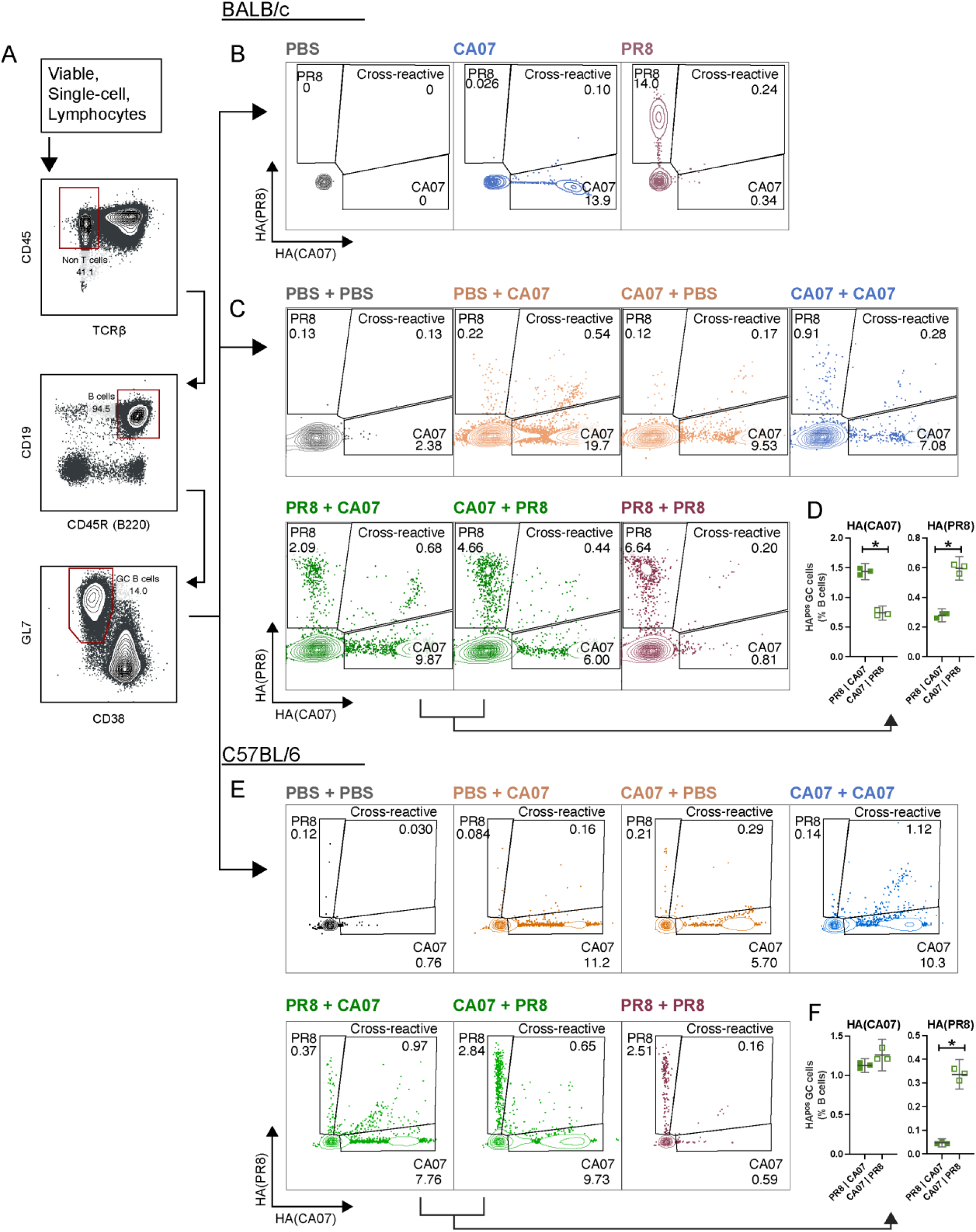
No imprinting effect on GC B cell specificity. **A)** GC B cells were first gated for live and single cells, and lymphocytes then gated as CD45^pos^ TCRb^neg^ CD19^pos^ B220^pos^ GL7^pos^ CD38^neg^ (see Extended data Fig. 3A). **B)** Representative plots of pooled mediastinal LN B cells harvested 4 weeks after a single infection of BALB/c mice with either CA07, PR8, or mock infected (n=5 per group). See Extended data Fig. 4A-B. **C-F)** Experimental setup depicted in Fig. 1B, with collection of lymphoid organs at day 56 after two subsequent influenza infections. All plots show representative replicates from triplicates performed on the pooled LN (see Extended data Fig. 4C-H). **C)** Pooled mediastinal LN were harvested from BALB/c mice (n=5 per group), infected as indicated. **D)** Direct comparison of HA specificities for GC B cells following heterologous influenza challenges of BALB/c mice with PR8+CA07 or CA07+PR8. Data shown as frequency of total B cell population. **E)** Pooled mediastinal LN from C57BL/6 mice (n=5 per group), infected as indicated. **F)** Direct comparison of HA specificities for GC B cells following heterologous influenza challenges of C57BL/6 mice with PR8+CA07 or CA07+PR8. Data shown as frequency of total B cell population. Statistical significance was calculated by Kruskall-Wallis’s one-way ANOVA, with a False Discovery Rate at Q=0.05 (D/F).

The same trend, with induction of both PR8 and CA07 specific B-cells after heterologous viral infections, was also seen in C57BL/6 mice (Fig. 2E, and Extended data Fig. 4, F-H). Of note, in BALB/c mice we observed higher numbers of GC B-cells against the strain used for secondary infections (Fig. 2D), but in C57BL/6 mice there were similar levels of CA07 specific cells in mice primed both with CA07 and PR8 (Fig. 2F).

### CA07 priming promote formation of MBC that are prone to differentiate into antibody secreting cells

The first influenza infection will produce plasma cells (PC) and MBCs. Different MBC subsets have different roles and functions with respect to how they respond to a secondary infection. More specifically, PD-L2^neg^CD80^neg^ MBC may efficiently seed new GCs, whereas PD-L2^pos^CD80^pos^ cells are more prone to differentiate into antibody secreting cells^13,23–25^. We therefore stained spleen derived HA^pos^ MBC (defined as CD38^pos^IgD^neg^) collected two weeks post a secondary infection for expression of PD-L2 and CD80 (Fig. 3A). All groups that had received a viral dose for the primary infection had 12-22% double positive (DP) (PD-L2^pos^CD80^pos^) HA-specific MBC, whereas the PBS+PBS and PBS+CA07 groups had 1-5% (p<0.001) (Fig. 3B). Thus, the first influenza exposure likely produced GC that lead to MBC expressing BCR with high specificity for its cognate antigen. The double negative (DN) MBC gave a different picture, and defined about 80% of MBC in the PBS+PBS and PBS+CA07 groups. Remaining groups had a clear decrease in the DN population (p<0.005), particularly pronounced for the CA07+PBS and CA07+PR8 groups. When comparing PR8+CA07 and CA07+PR8 mice we observed that the CA07 primed mice had significantly more DP MBC (p=0.036) as compared to the PR8 primed mice, indicating a reduced predisposition for future GC seeding in CA07 imprinted mice over PR8 imprinted mice (Fig. 3B).

**Figure 3:**
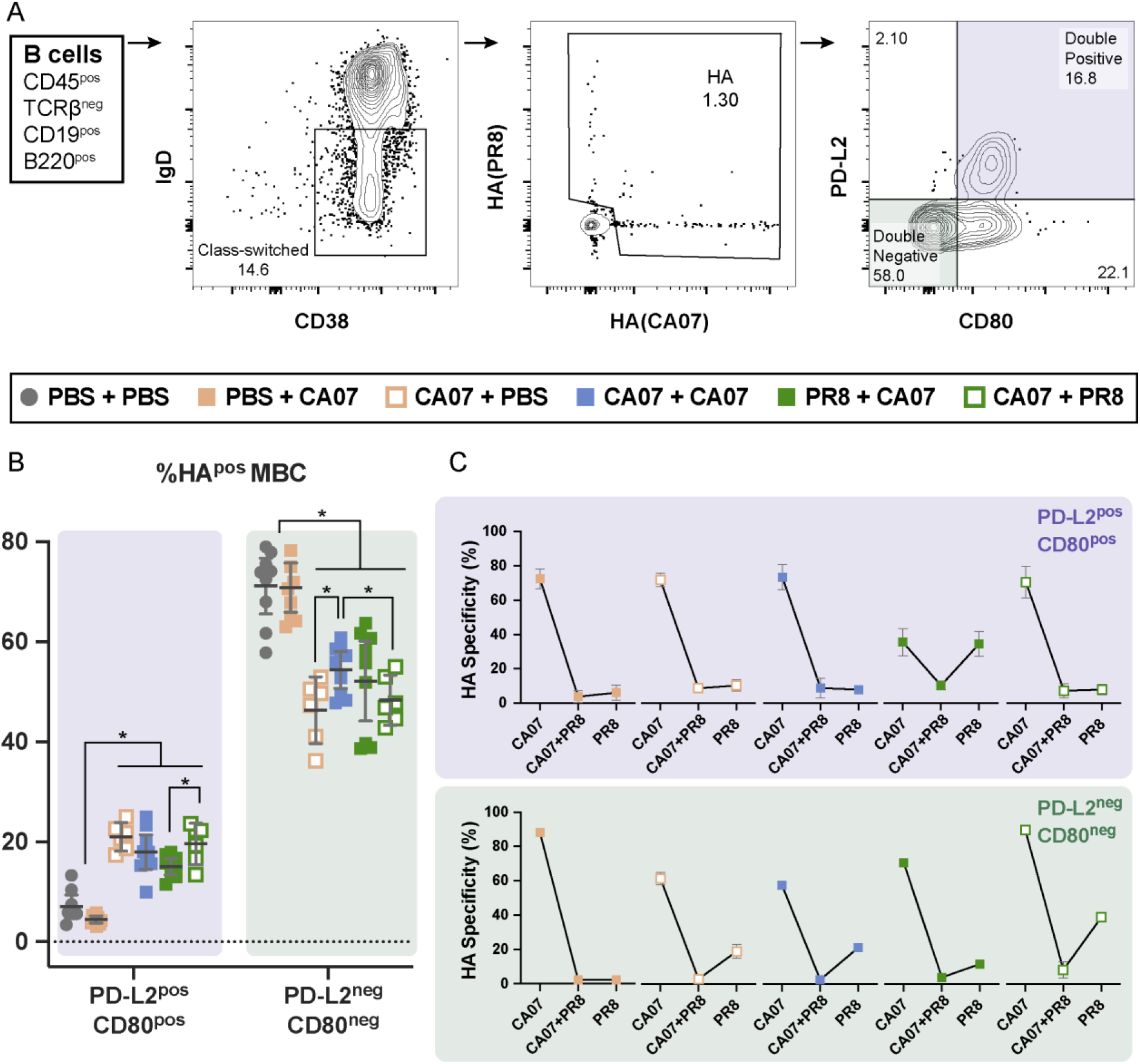
CA07 infection preferentially induced MBC prone to antibody secretion, in contrast to PR8. **A)** Gating strategy. Class-switched B cells (CD45^pos^ TCRβ^neg^ CD19^pos^ B220^pos^ IgD^neg^ CD38^pos^) were gated for HA specificity, and examined for CD80 and PD-L2 expression. **B)** Quantification of HA-specific MBC based on CD80 and PD-L2 expression in splenocytes collected 2 weeks post secondary viral exposure with either CA07, PR8, or PBS, as indicated. (Data was pooled from Extended data Fig. 1, experiments A/C/F, n=3×3 technical repeats from pooled groups). **C)** MBC that were either DP or DN for CD80 and PD-L2 were examined for specificity against HA(CA07), HA(PR8), or cross-reactivity against both. (Data from Extended data Fig. 1 experiment F, n=3 replicates per group). Statistical significance was calculated by Brown-Forsythe and Welch one-way ANOVA was used with a False Discovery Rate at Q=0.05 (B).

Separating the DP and DN MBC specificity for either PR8 and CA07 to quantify differences between priming with either CA07 and PR8, we found that about equal levels of DP MBC specific for either PR8 or CA07 were induced in the PR8+CA07 group (∼35% each). By contrast, the DP MBC induced in the CA07+PR8 group were almost exclusively specific for CA07 (70% CA07 compared to 7.9% PR8) (Fig. 3C), reinforcing the dominant imprinting effect following a primary infection with CA07 as compared to PR8.

### Phenotypes and mutations of HA binding B cells

To achieve a more detailed understanding of the impact sequential influenza exposures had on B cell responses, we next evaluated HA-binding B cells at the single-cell level. In brief, B-cells from draining LN in mice that 14 days earlier had been re-infected with either CA07 or PR8 were sorted and single cell RNA (scRNA) sequenced for gene expression (GEX), V(D)J, and cell surface protein libraries. The total B cell population in the PBS+PBS group was used as a control (Extended data Fig. 5).

GEX analyses demonstrated the presence of four or five main clusters of HA-binding cells, depending on the experiment (Fig. 4A-B). The largest cluster contained GC B cells that expressed *Bcl6* and *Aicda* (AID), for example, whereas “Circulating B cells” expressed markers indicative of either naïve B cells or MBC. The cluster “Pre-MBC” was characterized by higher expression of *Bach2* and other markers indicating GC BC predestined to become MBC^26^. Finally, a small cluster of plasma blasts (PB) with high levels of *Sdc1* (CD138) and *Prdm1* (BLIMP-1) were also present among the HA-binding B cells (Extended data Fig. 6). By using antibody-derived tag (ADT) based clustering from the surface protein library, the pooled cells could also be demultiplexed into different infection groups (see Extended data Fig.s 6-8 for results from similar and independent experiments). Looking at the phenotypes of the B cells in the different infection groups, we could see that the PBS+PBS cells were mostly naïve, expressing non-GC markers as well as IgD and IgM. The other infection groups contained mostly class-switched GC B-cells, with the exception of CA07+PR8 that expressed more class-switched circulating B cells than the other infection groups, and inverse with PR8+CA07 in particular (Fig. 4C).

**Figure 4:**
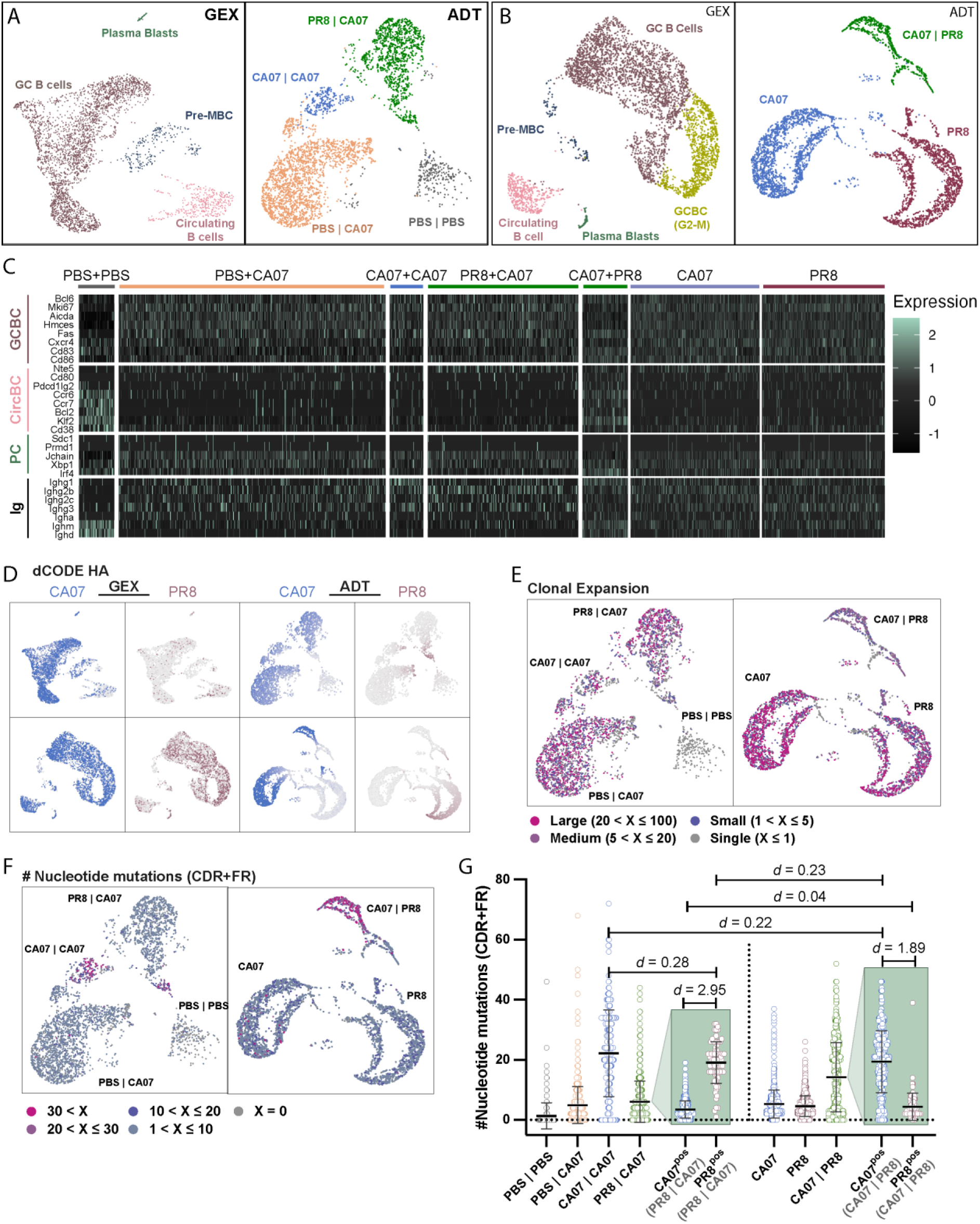
Single-cell RNA sequencing of HA specific B cells. Draining LNs were isolated from mice infected with either PBS+PBS, PBS+CA07, CA07+CA07, PR8+CA07, CA07+PR8, CA07+PBS, or PR8+PBS (n=8 per group), pooled within groups, and HA specific B cells isolated for BCR sequencing. **A-B)** Gene expression (GEX, B-cell subset, left) and antibody derived tag (ADT, included infection groups, right) based clustering of cells from the single cell RNA sequencing of HA specific B cells from two separate experiments (Extended data Fig. 5A-B). **C)** Heatmap of normalized expression of selected genes exemplifying the different clusters and immunoglobulin subtype for each infection group in A-B. **D)** B cell binding to HA(CA07) and HA(PR8) overlaid on GEX (left) and ADT (right) clustering. **E)** Clonal expansion by total number of cells in each clonotype, highlighted on ADT clusters, corresponding to the experiment depicted in A (left) and B (right). A clonotype was considered highly expanded if there were more than 20 cells (Large), moderately expanded if there were more than five cells, and lowly expanded if there were less than five but more than one cell (Small). **F)** Total number of nucleotide mutations of the BCR variable region sequence highlighted on ADT clusters, corresponding to the experiment depicted in A (left) and B (right). **G)** Quantification of mutations from F by infection groups. In addition, the HA(CA07) and HA(PR8) binding cells from PR8+CA07 and CA07+PR8 mice were separated and plotted (Green box). Differences in means were determined by Cohen’s d test where *d* = number of standard deviations.

Most of the isolated cells bound HA(CA07). HA(PR8) binding cells were also found in all groups infected with either virus, and in all B cell clusters, but mainly in mice solely infected with PR8 (Fig. 4D). This was in accordance with the above observations of cross-reactive responses following sequential exposure to CA07 (Fig. 1). When overlaying the BCR clonality information of each cell on the UMAP, we observed clonally expanded B cells in all infected groups, but not in the PBS+PBS control group (Fig. 4E). Using the total number of nucleotide mutations as an indication of time spent in GC reactions, we observed the highest rates of SHM in mice sequentially exposed twice to virus, either CA07+CA07, PR8+CA07, or CA07+PR8 (Fig. 4F). Quantifying the mutation counts for the different infection groups confirmed that cells originating from CA07+CA07 exposure were highly mutated, despite being a comparably smaller cluster (Fig. 4G). Looking specifically at the HA(PR8) specific cells from PR8+CA07 infected mice and the HA(CA07) specific cells from CA07+PR8 infected mice, we observed that they had similar mutation counts as compared to cells from CA07+CA07 infected mice (*d* = 0.28, *d* = 0.22 respectively) as well as each other (*d* = 0.23). As such, this indicates that they may have been maintained from the priming infection and undergone a new round of GC selection in parallel with potential de novo GC formation following the secondary infection.

In sum, we observed that the different HA-specific B cell subsets displayed a mutation rate corresponding to their expected phenotype, and that the second infection likely initiated new GC formation also as a secondary response (Fig. 4). This is in accordance with the previous observations regarding GC B cell distribution (Fig. 2).

### HA binding V(D)J sequences are diverse and do not rely on conserved motifs

To examine if the repeated exposures to influenza infection would lead to distinct motifs or patterns in the BCR of isolated HA specific B cells, we investigated sequence characteristics for the different infection groups. First, a network analysis of the HA-specific complementary determining region (CDR) heavy chain loop 3 (H3) clonotypes after different sequential exposures to the selected influenza H1N1 variants or PBS demonstrated that the uninfected mice displayed very limited connectiveness. This indicated that very little directional affinity maturation had taken place, as expected (Fig. 5A-B). The mice exposed to PBS+CA07 displayed some connectivity, but markedly less as compared with the CA07+CA07, PR8+CA07, and CA07+PR8 groups. The CA07+CA07 group displayed the largest clusters of similar clonotypes, whereas the PR8+CA07 and CA07+PR8 groups were a bit more diverse. However, this did not translate into clearly different conserved CDRH3 motifs between HA-specificities. The most noteworthy difference was the enrichment of asparagine in position four for HA(CA07)-specific sequences, as this was not observed for either HA(PR8) or cross-reactive sequences (Fig. 5C-D). Unfortunately, the low number of cross-reactive clones made CDR3 sequence analysis unreliable.

**Figure 5:**
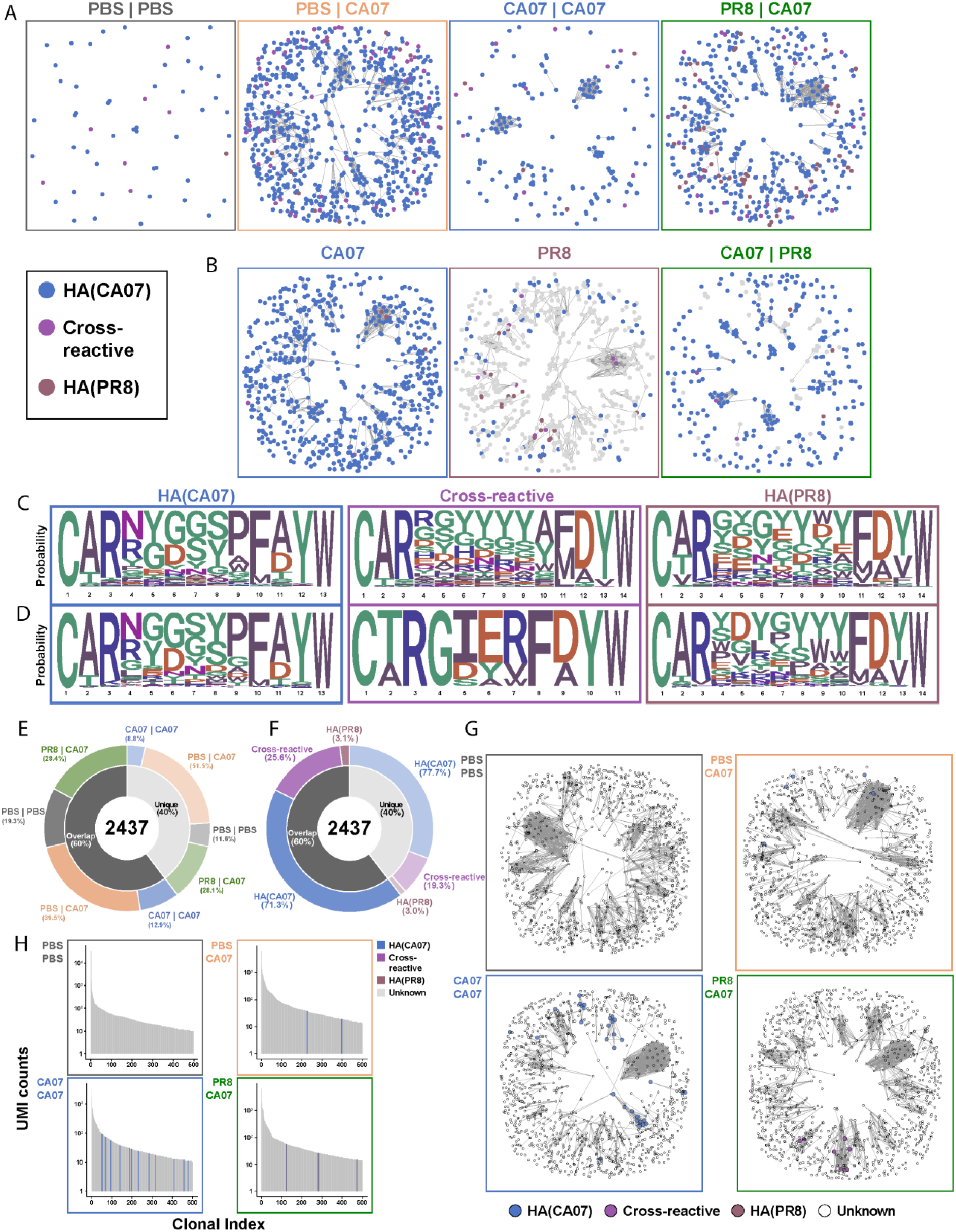
BCR repertoire analysis revealed a diverse repertoire of HA-binding sequences. A-B) Network analysis of HA-specific CDRH3 sequences from draining LNs (**A**: mice from Extended data Fig. 5A, B: mice from Extended data Fig. 5B). Two CDHR3 sequences are joined by a line if only a single amino acid mutation differ between them. **C)** Logoplots of HA(CA07)-specific, HA(PR8)-specific, and cross-reactive CDHR3 sequences, based on sequences from three merged sequencing runs described in Extended data Fig. 5A, C, and D. **D)** Logoplots of HA(CA07)-specific, HA(PR8)-specific, and cross-reactive CDHR3 sequences, based on one sequencing run described in Extended data Fig. 5B. **E)** Circos plot of infection group distribution between CDHR3 sequences unique to single cell LN sequences, or sequences overlapping between single cell LN sequences and bulk PC sequences. **F)** Corresponding circos plot of HA-specificity for unique and overlapping sequences. **G)** Representative bulk PC CDRH3 networks of the 1000 most prevalent clones for each infection group. Highlighted sequences are shared clonotypes with single-cell LN sequences, and HA-specificity is annotated accordingly. **H)** Top 500 expanded bulk PC clonotypes from representative mice of each infection group. Highlighted clonotypes are shared with single-cell LN sequences and HA-specificity annotated accordingly. Fig. E-H are data from experiment outlined in Extended data Fig. 5A.

PCs are the main producers of serum IgG. To better link the serum antibody responses (Fig. 1) to the single cell BCR sequences, we performed bulk V(D)J sequencing of the PC compartment from a selection of the same mice that we had also collected HA-specific B cells from. Only cells from PBS+PBS, PBS+CA07, CA07+CA07, and PR8+CA07 were available for this analysis. In brief, we isolated PCs from the bone marrow of each mouse individually and extracting the RNA for BCR sequencing. Unfortunately, this did not allow for labelling of HA-specific sequences, but by comparing sequences found in both compartments we were able to identify HA-specific PC clonotypes. Firstly, we found that 60% of HA-specific LN sequences were also found among the PC clonotypes, while 40% were unique to the LNs. Secondly, we observed that the distribution between infection groups remains similar for both overlapping and unique LN sequences, indicating that the infection did not skew the sampling (Fig. 5E). The same was seen for HA-specificity, where no large differences were observed between the overlapping and unique sequences (Fig. 5F). This suggests that limited sampling of LN B cells was the main reason for this low level of overlap, and that if more cells had been sequenced this overlap would increase.

Next, we examined the top 1000 most expanded PC clones for each mouse and found that we could recover HA-specific sequences among them. Sequence-similarity-based network analysis revealed connected clusters of clones where one or more cells were recovered among the HA-specific LN sequences (Fig. 5G). This indicated that the remaining cells in these clusters were also likely to be HA-specific. By then looking at the top 500 most abundant bone marrow PC clones, we found shared clones in infected mice but not in non-infected control mice (Fig. 5H). As such, we have demonstrated that the primary infection could be clearly seen in the general PC compartment, and that relatedness increased with subsequent infections.

### In vitro validation of scRNA-sequenced clones confirms specificity and show varying neutralization capabilities

As a control of assigned sequence specificities (Fig.s 4-5), we wanted to validate a selection of the obtained BCR sequences. Thus, the paired heavy chain (H) and light chain (L) sequences from eleven clonotypes were selected from the pool of assigned HA binding sequences for recombinant expression (Fig. 6A-B, Extended data table 1). The sequences were selected to cover a range of binding specificities, including CA07 only, PR8 only, and cross-reactivity for both CA07 and PR8. Further, the sequences were selected to cover a wide range of clonal expansions and mutation rates. Lastly, the sequences were also selected from multiple independent sequencing runs (Extended data Fig. 1, experiments D and E, extended data Fig. 5A, C).

**Figure 6:**
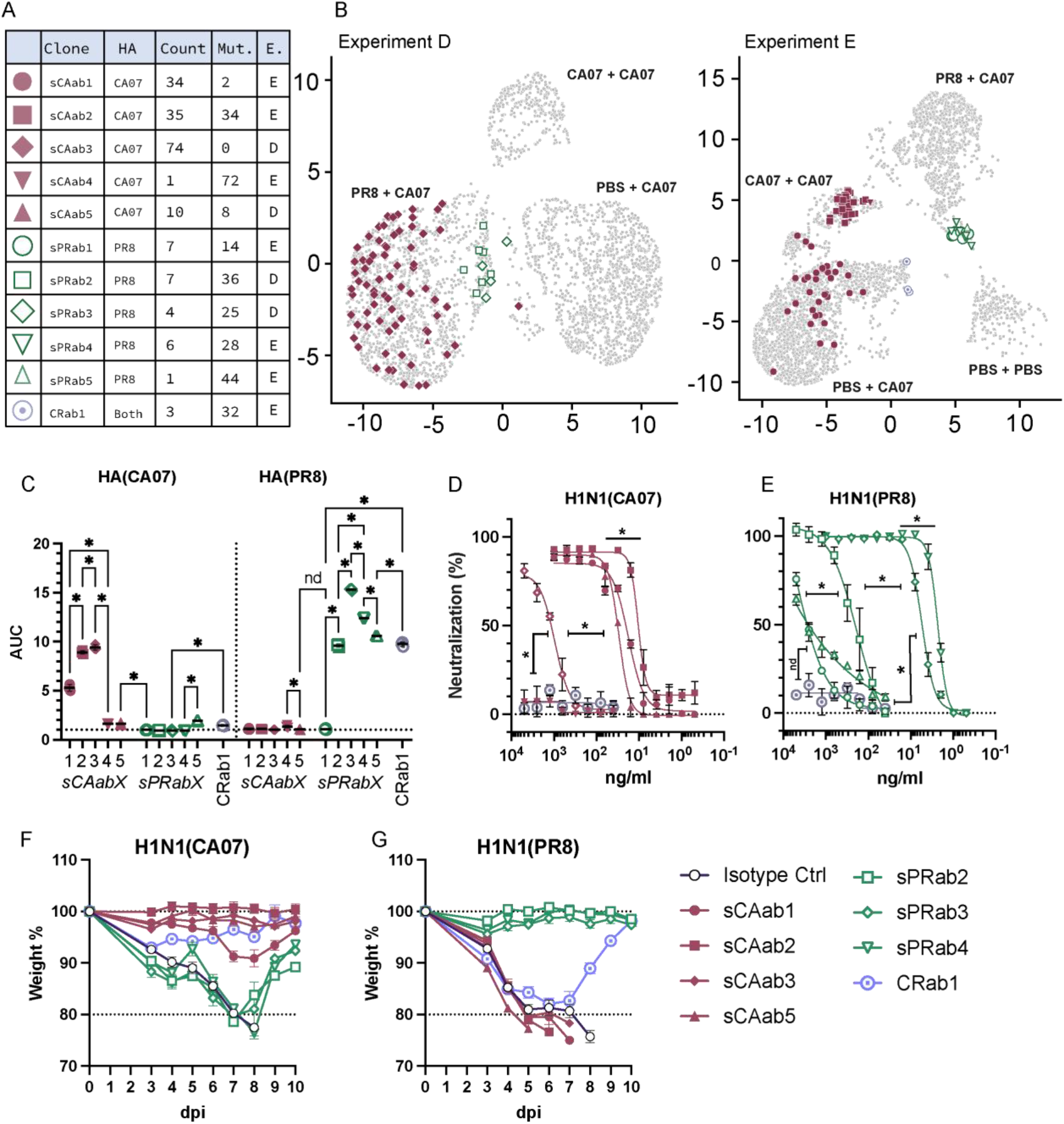
Experimental validation of in silico assignment of HA sequence specificity. **A)** Summary of selected in silico defined clones from experiments D and E (Extended data Fig. 1 and Extended data Fig. 5A&C). Depicted are their assigned HA binding from the scRNA sequencing, the number of cells detected for each clonotype (Count), the total number of mutations observed in each clonotype (FR and CDR regions combined) (Mut.), and which experiment (E.) they were extracted from in Extended data Fig. 1 and Extended data Fig. 5A&C. **B)** Selected clones highlighted on ADT-based UMAP clustering for Exp D and E. **C)** Binding of purified mAbs to HA(CA07) (left) and HA(PR8) (right) (n=3 independent replicates). Statistical significance was calculated with Brown-Forsythe and Welch one-way ANOVA (FDR Q=0.05). **D-E)** In vitro neutralizing potential of mAbs against CA07 and PR8 (n=3). Calculated IC_50_ concentrations were used for assessment of statistical significance by Brown-Forsythe and Welch one-way ANOVA (FDR Q=0.05). **F**) In vivo protection of mAbs against a lethal influenza challenge 2xLD50 with H1N1(CA07). Data merged from three separate experiments, n=17 for Isotype control, n=8 for sCAab1, sCAab3, sCAab5, sPRab2, sPRab3, sPRab4 and CRab1, n= 12 for sCAab2. **G**) In vivo protection of mAbs against a lethal influenza challenge 2xLD50 with H1N1(PR8). Data merged from three separate experiments, n=20 for Isotype control, n=8 for sCAab1, sCAab2, sCAab3, sCAab5, sPRab2, sPRab4 and CRab1, n= 12 for sPRab3.

The selected clones were expressed as monoclonal human IgG1, purified, and tested for binding to HA. Importantly, all HA(CA07) assigned binders bound HA(CA07) and 4/5 HA(PR8) assigned binders bound HA(PR8), although at varying degrees. The cross-reactive clone successfully bound HA(CA07) and HA(PR8) (Fig. 6C). Furthermore, we found that one HA(PR8) assigned clone bound HA(CA07) (p=0.003), and that one HA(CA07) assigned clone had cross-reactivity also for HA(PR8) (p=0.044).

We next evaluated the neutralization capabilities of the produced mAbs against the corresponding influenza virus *in vitro*. Here, 4/5 HA(CA07) specific mAbs neutralized CA07 (Fig. 6D). The same 4/5 HA(PR8) specific mAbs that bound HA(PR8) also neutralized PR8, but with varying potencies (Fig. 6E). The cross-reactive mAb was not capable of neutralizing either virus, which could be due to specificity for an epitope outside of the globular head^27,28^.

We then further verified the specificity of the produced mAbs by seeing whether they could protect mice *in vivo* against a lethal influenza virus infection. Here, all CA07 specific mAbs, as well as the cross-reactive mAb, provided complete protection against CA07. The mice receiving PR8 specific mAbs lost weight to the same extent as the isotype control mice, but one mouse in each group stabilized above the threshold for euthanasia and recovered (Fig. 6F). All PR8 specific mAbs provided complete protection against PR8. The cross-reactive mAb provided partial protection with a significant weight loss, with 3/8 mice surviving. Neither the CA07 specific mAbs nor the isotype control provided any protection against PR8 virus (Fig. 6G).

In sum, we were able to verify that the HA specificity assigned during B cell isolation and BCR sequencing corresponded well to functional HA binding and protection against disease. However, the experimentally observed neutralizing capacity of the selected recombinant mAbs varied, which is expected in the absence of information on epitope specificity.

## Discussion

We here explored the ramifications of immune imprinting not only on the serum antibody responses, but also on the cellular composition and diversity of the immune repertoire following two sequential viral challenges with influenza PR8 and CA07 viruses. Intriguingly, mice primed with CA07 manifested attenuated serological responses to PR8 upon a heterologous re-infection, while the inverse scenario did not elicit a similar outcome. This imprinting effect was not observed at the B cell level, where we found a wide range of HA-binding BCR sequences devoid of conserved binding motifs and encompassing a broad mutational spectrum. Clonal analyses demonstrated the existence of overlapping antigen-specific clonotypes within both lymph nodes and bone marrow. The subsequent reconstitution of selected clonotypes as human IgG1 antibodies lent credence to our sequence data interpretations, and provided insights into the neutralizing capabilities of the cloned sequences.

The degree of similarity between the antigens likely contributed to the observed imprinting. The HA of CA07 and PR8 share an approximate 82% amino acid identity, which is a larger distance than that typically witnessed during annual seasonal drift ^29^. As a reference, the WHO will typically update vaccine recommendations when discrepancies between circulating and vaccine-incorporated strains exceed 3% ^30^. However, the ∼18% disparity between CA07 and PR8 is not unrealistic, corresponding approximately to the evolutionary breach associated with the emergence of the 2009 H1N1 (“swine flu”) pandemic^30^.

In our observations, primary exposure to the CA07 viral strain led to an upsurge in PD-L2^pos^CD80^pos^ HA-specific MBCs, paired with a diminishing of PD-L2^neg^CD80^neg^ MBCs, a phenomenon observed to a lesser degree following PR8 exposure. PD-L2 and CD80 are both B cell markers acquired during T-cell dependent GC reactions^31^, and PD-L2^pos^CD80^pos^ has previously been shown to guide memory responses towards antibody secretion rather than GC reentry ^13,25,32^. In addition, it has been shown that PD-L2^pos^ MBCs and CD80^pos^ MBCs remain in the lung after an influenza infection, and rapidly differentiates into Ig-secreting PC^33^. The observed increase in PD-L2^neg^CD80^neg^ MBCs after PR8 imprinting could explain the heightened HA(CA07)-specific responses upon secondary CA07 infection, while the higher number of PD-L2^pos^CD80^pos^ MBCs following a priming event with CA07 correspondingly could lead to a preferred activation of pre-existing MBCs. While selected BCR sequencing of these distinct MBC subsets was not performed in our study, we noted that, post reinfection with PR8, almost none of the CA07-primed PD-L2^pos^CD80^pos^ MBCs bound HA(PR8). This contrasts with MBCs primed by PR8 and re-infected with CA07, and where roughly equal binding to both HA antigens were detected. Such findings underscore that initial exposure to PR8 primes B cells for response versatility against CA07, an assertion further supported by the variation in mutation rates we documented (Fig. 4G).

A six-week interval was implemented between primary and secondary infections, exceeding the routine prime-boost schedule for de novo vaccinations against respiratory viruses ^34,35^. The type of vaccination will influence GC reaction duration, ranging from several weeks and to over half a year^36–38^. A longer interval between prime and boost vaccinations can induce a stronger neutralizing antibodies response and better long-term protection^39–41^. Viral infections often induce long-lived GC reactions, but these are not maintained solely by founding virus-specific cells^42^. This could affect subsequent immune responses as there may be a limited number of concurrent GC reactions permitted in one LN ^43^.

At day14 after the second influenza exposure, we identified a reduction of GC B cells in all mice that had received the same strain during imprinting and re-infection, but that was not observed for the CA07+PBS C57BL/6 mice (Extended data Fig. 4). This could be attributed to a combination of ongoing GC reactions and circulating serum antibodies preventing further GC activation by means of epitope masking ^44–47^. To mitigate this confounding factor, it would be necessary to allow the GC reactions to subside and allow for a more direct study of the MBCs that remained in the mice.

To validate the assignment of HA specificity to the sequenced antigen specific BCR repertoire, and gain insight into the relationship between mutational load and neutralization capacities, we selected 11 BCR sequences representative of a wide array of clonal expansions and mutational rates for the generation of corresponding mAbs. Highly mutated BCRs are often associated with high affinities and/or broad specificity^48^. However, two of our mAbs (SCA2 and SCA3) exhibited similar neutralization capabilities despite SCA3’s germline status and SCA2’s high mutational load. Furthermore, despite being the most highly mutated unique clone, SCA4 failed to neutralize the homologous virus. This provides further backing to the notion that certain germline sequences are in an advantageous position for binding viral antigens^49–53^.

In conclusion, our study revealed clear differences in the immune response to secondary infections following a first priming with either PR8 or CA07. The imprinting dominance translates into clear effects on serum antibodies, but smaller differences on B cell specificities.

## Material and methods

### Animals

Female BALB/c or C57BL6 mice (Janvier) at 6-8 weeks of age were used for all experiments. All animals were housed at the Department of Comparative Medicine, Oslo University Hospital, Oslo, Norway, under minimal disease conditions until viral infection, at which point the mice were housed in a BSL2 classified unit approved for work with infectious agents. All experiments were approved by the Norwegian Food Safety Authority (FOTS id: 20892 and 27552).

### Influenza challenge and vaccination of mice

Female BALB/c mice were infected intra nasally (i.n.) with 0.25×LD_50_ for sublethal infections, or 2xLD_50_ for lethal infections, with either A/California/07/2009(H1N1) (CA07) or A/Puerto Rico/8/1934(H1N1) (PR8). For vaccinations, 50 µg DNA encoding APC-targeted HA ^19,20^ was injected intra muscularly (i.m.) into quadriceps followed by electroporation of the injection site.

Blood samples were obtained every other week by saphenous vein puncture (<80µl), or during harvesting of lymphoid tissues via cardiac puncture and terminal bleeding (>500µl). For serum transfer experiments, 200µl pooled serum was injected intravenously 24 hours prior to infection.

### Enzyme-linked immunosorbent assay (ELISA)

Microtiter plates, 96-wells, were adsorbed with 0.5 µg/mL recombinant HA protein (produced in-house, or Cat. No. 11085-V08H, 11684-V08H, 11055-V08H4, 11684-V08H1, SinoBiological) at 4°C for 18 hours before being blocked by 1% BSA in PBS at room temperature for 60 minutes. Serum from immunized mice were serially diluted 3-fold in 0.1% BSA and 0.05% Tween20 in PBS from a 1:50 starting dilution, and incubated at 4°C for 18 hours. Serum IgG was detected by anti-mouse IgG-HRP (1:5000, Cat. No. A2554. Merck) at room temperature for 60 minutes. Mouse IgG1 was detected by anti-mouse IgG1-HRP (1:10 000, Cat. No. ab97240. Abcam) and mouse IgG2a was detected by first adding biotinylated anti-mouse IgG2a[a] (1:500, Cat. No. 553502. BD Pharmingen) before addition of Streptavidin-HRP (1:3000, Cat. No. 7100-05. Southern Biotech). Next, plates were developed with TMB solution (Cat. No. CL07-1000ML, Merck or Cat. No. 2023, Thermofisher) for 10 minutes before quenching by 1M H_2_SO_4_. Absorbance was measured at 450 nm using a Wallac Envision multiplate reader (PerkinElmer). Area under the curve (AUC) was calculated by the trapezoidal method 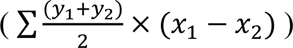 with Log_10_(dilution factor) as x-values and absorbance as y-values.

### Competition ELISA

Serum was diluted to the dilution that gave OD450 ≈ 1 in ELISA and incubated with 10µg/ml recombinant HA from either CA07 or PR8 (produced in-house) and incubated for 120 minutes at room temperature. Control consisting of serum without HA was performed in parallel. Microtiter plates, 96-wells were adsorbed with recombinant HA and blocked in the same way as described above. The premixed serum+HA was then added to the microtiter plates, incubated and detected with anti-mouse IgG-HRP as described above. Signal was detected using TMB substrate as described above and inhibition was determined as loss of signal compared with control serum without HA pre-adsorption.

### Microneutralization assay

Sera from individual mice were mixed with Receptor Destroying Enzyme (RDE) (Cat. No. C8772-1VL. Merck) at a ratio of 1:3 and incubated at 37°C for 18 hours before being inactivated at 56°C for 30 minutes. A two-fold serial dilution of serum:RDE was mixed with virus PR8 or CA07 at 25×TCID_50_ (final concentration), and pre-incubated at 37°C for 120 minutes before added to plates with 2×10^4^ MDCK cells per well. The plates were then incubated for 20 hours in a humidified, 5% CO_2_ incubator at 37°C. The cells were next washed with PBS and fixed with ice-cold 80% acetone in PBS for 12 minutes, and left to air dry. Next, plates were incubated with 1% BSA for 60 minutes and washed (0.05% Tween20, PBS), and 1 µg/ml biotinylated anti-NP (in-house, clone H16-L10-4R5) was added for 60 minutes. A secondary incubation of Streptavidin-HRP (1:3000, Cat. No. 7100-05. Southern Biotech), or anti-mouse IgG-HRP (1:3000, Cat. No. A2554. Merck) for 60 minutes was performed before development by TMB solution (Cat. No. CL07-1000ML, Merck or Cat. No. 2023, ThermoFisher) and 1M H_2_SO_4_, as described above.

Microneutralization of mAbs were performed as above, except that mAbs were added instead of serum at 2µg/ml, 5µg/ml or 10µg/ml starting concentrations. No RDE treatment or heat inactivation was used for the mAbs.

### Flow cytometry analysis

Single cell suspensions were made from mouse tissues by manual passage through a 70 µm cell strainer (Cat. No. 732-2758. VWR). Splenocytes and LN cells were next treated with RBC lysis buffer (In-House: 150mM NH_4_Cl, 10mM Na_2_CO, 0.1mM Na_2_EDTA, pH 7.4) for 10 minutes on ice. Cells were next washed with PBS and viability stained with GhostDye510 (1:400. Cat No. 13-0870-T500. Tonbo Bioscience), blocked with 25% normal rat serum (Cat No. 10710C. ThermoFisher), before being stained for the following surface markers: Memory B cell analysis: TCRb-BB700 (1:200. Cat No. 745846. BD Bioscience), CD19-APC/Cy7 (1:200. Cat No. 557655. BD Bioscience), CD38-BUV395 (1:200. Cat No. 740245. BD Bioscience), CD45-BUV496 (1:200. Cat No. 749889. BD Bioscience), CD45R(B220)-BUV805 (1:200. Cat No. 748867. BD Bioscience), CD73-BV711 (1:200. Cat No. 752736. BD Bioscience), CD80-PE/Cy7 (1:200. Cat No. 104734. Biolegend), CD273(PD-L2)-BV421 (1:200. Cat No. 564245. BD Bioscience), IgG1-BV605 (1:100-200. Cat No. 563885. BD Bioscience), IgG2a-BV605 (1:100-200. Cat No. 564024. BD Bioscience), IgM-BUV737 (1:200. Cat No. 749079. BD Bioscience), IgD-BV786 (1:200. Cat No. 563618. BD Bioscience), and Avitag®-biotinylated (Cat. No. MSPP-BIRA500. Avidity Biosciences) and PE/APC-Klickmer®-coupled HA (10 µl 32 nM per sample). GC B cell Analysis: TCRb-BB700 (1:200. Cat No. 745846. BD Bioscience), CD19-APC/Cy7 (1:200. Cat No. 557655. BD Bioscience), CD38-BUV395 (1:200. Cat No. 740245. BD Bioscience), CD45-BUV496 (1:200. Cat No. 749889. BD Bioscience), or CD45.1-Alexa Fluor® 700 (1:200. Cat No. 110724. Biolegend) plus CD45.2-BV711 (1:200. Cat No. 109847. Biolegend), CD45R(B220)-BUV805 (1:200. Cat No. 748867. BD Bioscience), CD273(PD-L2)-BV421 (1:200. Cat No. 564245. BD Bioscience), IgG1-BV605 (1:100-200. Cat No. 563885. BD Bioscience), IgG2a-BV605 (1:100-200. Cat No. 564024. BD Bioscience IgM-BUV737 (1:200. Cat No. 749079. BD Bioscience), IgD-BV786 (1:200. Cat No. 563618. BD Bioscience), GL7-Alexa Fluor® 488 (1:200. Cat No. 144612. Biolegend), and Avitag®-biotinylated (Cat. No. MSPP-BIRA500. Avidity Biosciences) and PE/APC-Klickmer®-coupled HA (10 µl 32 nM per sample, Cat. No. DX01K. Immudex). All staining buffers included BD Horizon™ Brilliant Stain Buffer (1:50. Cat No. 563794). Samples were analyzed on a FACSymphony A5 (BD Bioscience) running FACSDiva (BD Bioscience), or an Aurora (Cytek) running SpectroFlo®. Subsequent analysis was performed in FlowJo v10 (BD).

### Flow cytometry cell sorting and single cell sequencing

Splenocytes and mediastinal LN cells were acquired as described above. MBC were first isolated by magnetic separation using the Memory B cell Isolation kit (Cat. No. 130-095-838, Miltenyi) according to the manufacturer’s instructions. LN B cells were isolated using EasySep™ Mouse Pan-B Cell Isolation kits (Cat. No. 19844. Stemcell Technologies) according to the manufacturer’s instructions. MBC and LN B cells were then viability stained with GhostDye (1:400), blocked with 25% normal rat serum, and stained with TCRb-BB700 (1:200. Cat No. 745846. BD Bioscience), CD19-APC/Cy7 (1:200. Cat No. 557655. BD Bioscience), B220-PE/Cy7 (1:200. Cat No. 60-0452-U100. Tonbo Bioscience) and CD38-PE/Dazzle™ 594 (1:200, Cat No. 102730. Biolegend). LNs were in addition stained with GL7-Alexa Fluor® 488 (1:200. Cat No. 144612. Biolegend) and MBC were stained with IgG-APC as part of the magnetic isolation protocol (Cat. No. 130-095-838, Miltenyi). The cells were also stained with dCODE® PE-Klickmer (Immudex) coupled CA07 HA, PR8 HA and OVA (15:1 Ag:Klickmer ratio) with three unique barcodes. Groups were tagged with unique 10X compatible barcode TotalSeq™ C hashtag Abs (C0301-8, Cat No. 155861, 155863, 155865, 155867, 155869, 155871, 155873, 155875. Biolegend).

For Experiments in Extended data Fig. 5A-C: Viable, antigen positive B cells (TCRb^neg^, CD19^pos^) were sorted using either a FACSaria II (BD bioscience), (A and C), or a FACSMelody (BD Bioscience) (B). For Experiments in Extended data Fig. 5D: GC B cells (CD19^pos^, B220^pos^, CD38^neg^, GL7^pos^) and MBC (CD19^pos^, B220^pos^, CD38^pos^, IgG^pos^) were sorted separately using a FACSaria II (BD bioscience).

Sorted cells were prepared for single cell RNA sequencing using 10X Genomics (Cat. No. PN-1000266, PN-1000267) libraries for gene expression (Cat. No. PN-1000196), V(D)J (Cat. No. PN-1000255), and surface tag (Cat. No. PN-1000541) sequencing according to the manufacturer’s instructions, with ∼40 000 reads per cell for RNA expression and ∼8000 reads per cell for BCR sequencing. Assistance with library construction was given by the Oslo University Hospital Genomics Core Facility. Sequencing was performed by Oslo University Hospital Genomics Core Facility and Norwegian Sequencing Center at the University of Oslo.

### Bone marrow preparation and bulk BCR sequencing

Bone marrow was collected and CD138 positive cells were isolated by magnetic separation according to the manufacturer’s instructions (Cat. No. 18957, Stemcell Technologies). Total RNA was then isolated (Cat. No. 74104, Qiagen) and libraries were prepared with the NEBNext® Immune Sequencing Kit according to the manufacturer’s instructions (Cat. No. E6330S, New England Biolabs). Finished libraries were sequenced on an Illumina Miseq v3 with 300bp paired-end reads. Sequencing was performed by the Norwegian Sequencing Center at the University of Oslo.

### Bulk and single-cell sequencing preprocessing

FASTQ files from bulk and single-cell sequencing were processed by MiXCR version 4.1.0 with UMI correction, using a BALB/c mice reference set from ORGDB^54^. In brief, each sequencing read was assembled into a consensus sequence with a minimum quantity of 1 to retain all data. UMI-corrected reads were then aligned to the BALB/c mice reference set to determine the gene features (V, D, J genes and CDRs). Then the reads were collapsed into clonotypes by the features of the V(D)J region and the C gene. 10X single-cell BCR sequencing data was also processed using MiXCR, using the preset 10x-vdj-bcr, with additional steps for partial assembly of CDR3 sequences, and assembling the longest possible contigs.

### Single-cell gene expression analysis

In brief, for the gene expression analysis and clustering, the output of “cellranger count” was processed using Seurat v4 ^55^. First, hashtag demultiplexing was done by assigning one hashtag sequence to each cell using the “HTOdemux” function from Seurat. Doublets were then filtered and removed using the “doubletfinder” package ^56^, after which the HA specificity was assigned. This was done by comparing the log-transformed number of HA(CA07) and HA(PR8) reads for each cell to threshold-values between 0.9-1.1 (based on the run). If different cells within a single clonotype were then assigned both cross-reactive and specific for only one HA, they were reassigned as specific for that single HA.

### Repertoire analysis

#### CDR3 sequence overlap

CDR3 sequence overlap between samples was measured by Jaccard’s overlap index, with values ranging from 0 (no overlap) to 1 (total overlap) and calculated using the following formula: J(A,B) = |A∩B| / |A∪B|, where A and B represent the CDR3 amino acid sequences of two repertoires.

#### CDR3 network analysis

Sequence similarity network was constructed using the CDR3 amino acid sequences of each sample. Briefly, a network was drawn using the R package *igraph*^57^ based on the whole repertoire (for single-cell sequencing samples) or the top 1000 clonotypes (for bulk sequencing samples). Within each network, a node is one unique CDR3 amino acid sequence and edges between nodes are drawn if the internode Levenshtein distance (LD) is at most one (one amino acid sequence change).

#### CDR3 sequence probability distribution

For all single-cell sequencing samples in each infection group, the probability distribution of residues for each position in the CDR3 sequence was visualized using the R package *Platypus*^58^. The number of residues chosen for each logo plot was based on the most frequent CDR3 length.

All code is available at: https://github.com/csi-greifflab/Preexisting-immunity-influenza-mice.

### Production of monoclonal Abs from single cell sequences

Paired H and L sequences from cells with interesting characteristics were ordered as human IgG1 constructs (Twist Biosciences). For each HC clonotype selected, the most frequent LC clonotype associated with that HC clonotype was chosen as the HC-LC pair for mAb production.

### ELISA validation of monoclonal Abs

96-well microtiter plates were adsorbed with 2 µg/mL, 1 µg/mL, or 0.5 µg/mL recombinant HA protein (Sino Biological, or produced in-house) at 4°C for 18 hours before being blocked by 1% BSA in PBS at room temperature for 60 minutes. After washing, mAbs were diluted to 5 µg/ml, 10 µg/ml, or 2.5 µg/ml and added to the first row before being diluted 3-fold and incubated at room temperature for 2 hours. Bound mAbs were then detected with HRP-coupled anti-human IgG1 (1:2000. Cat. No. MBS675124, MyBioSource) for 60 minutes before being developed with 100 µl TMB (Cat. No. CL07-1000ML, Merck or Cat. No. 2023, Thermofisher) for 5 minutes and quenched as previously described. Absorbance was measured, and AUC was calculated as described above.

### Statistical analysis

For analyses where individual mice were treated separately, such as serum ELISA and neutralization analysis, non-parametric tests were performed. For tests where mice were pooled beforehand the parametric test versions were allowed. For serum ELISA analysis, each timepoint was compared separately by a Kruskall-Wallis one-way ANOVA correcting for multiple comparisons by False Discovery Rate (FDR) using the two-stage Benjamini, Krieger and Yekutieli step-up method with a threshold Q-value of 0.05. Similar Kruskall-Wallis tests with FDR adjustments were used for IC_50_-values. Mann-Whitney tests were instead performed when only two groups were included, as in the competition ELISA.

For flow cytometry analysis, Brown-Forsythe and Welch one-way ANOVA was used with the same FDR as above. Mann-Whitney tests and Welch’s t-test were instead used for non-parametric and parametric analysis respectively when only two groups were included.

Figures are indicated with a discovery, “*”, while the adjusted p value is described in the text. For mutational load analysis, Cohen’s d test were used to determine effect size according to |Mean_1_ - Mean_2_|/SD_Pool_, where 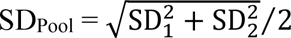. This gives a difference of means, “*d”*, in standard deviations.

All statistical analysis was performed in Graphpad Prism v10, except Cohen’s d test which was performed in Microsoft Excel.

## Acknowledgement

We wish to thank the Genomics Core Facility at Oslo University Hospital for performing the library preparations and next-generation sequencing, and also the Norwegian Sequencing Center at the University of Oslo for performing parts of the sequencing. We also wish to thank the Department of Comparative Medicine at Oslo University Hospital for maintaining the mouse colony used for these studies.

## Author contributions

ET, VG, and GG designed the study. ET, TMK, and MLTD performed the experiments, and ET and MLTD analyzed the mouse experiments. ET and QKL performed the bioinformatic analysis. QKL and VG developed bioinformatics tools for the analysis. ET and GG wrote the manuscript. All authors contributed with comments and feedback on the manuscript.

## Data availability

Data from experiments with flow cytometry and mouse experiments are deposited in the ImmPort repository. Authors can also be contacted directly for sharing of data.

