## Extended Data Figures 1 to 8, and Extended Data Table 1 for "Influenza H1N1 Imprinting Uncouples Serological and B-Cell Responses in Mice"

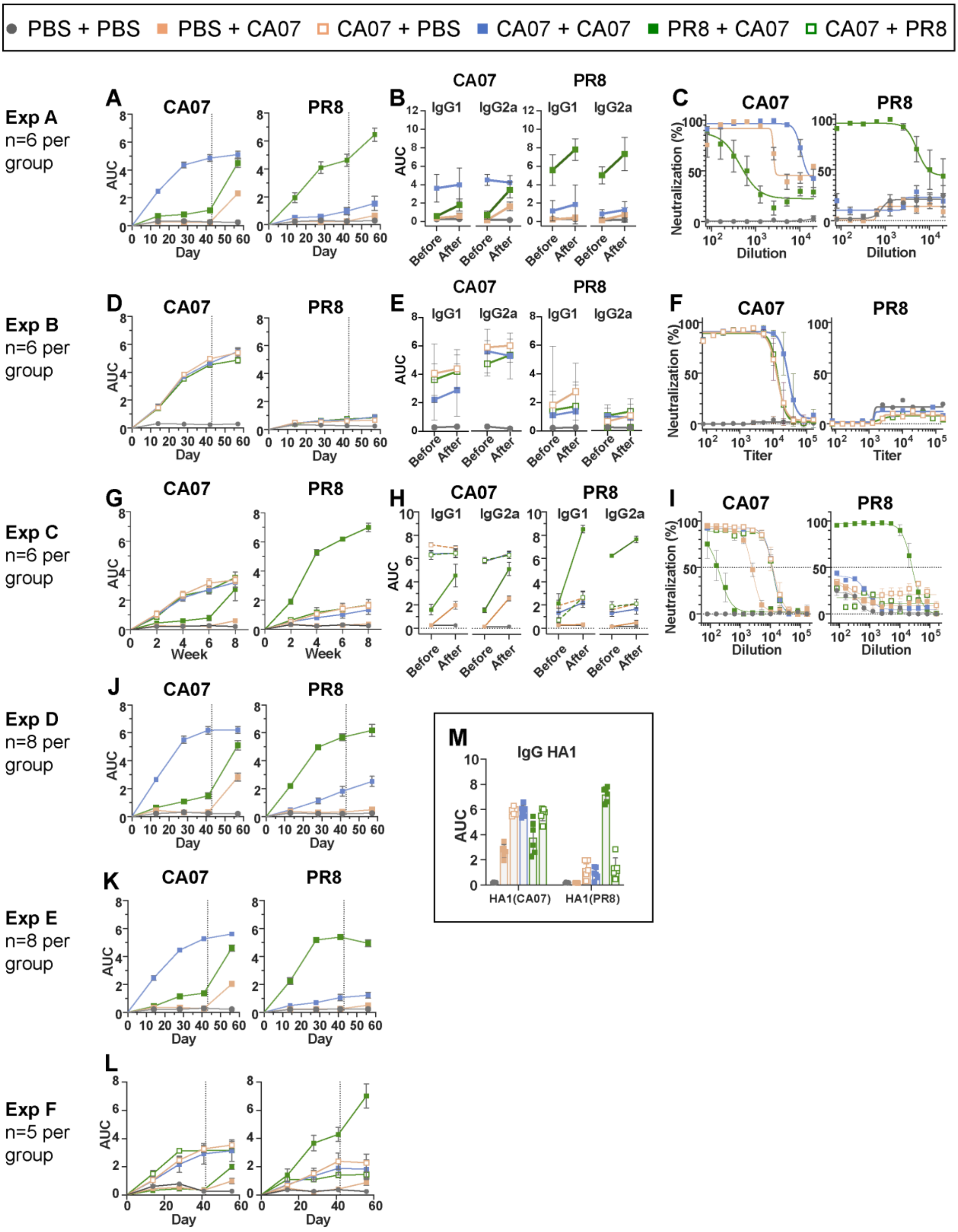


**Extended data figure 1: Antibody responses from 6 independent experiments in BALB/C (A-K+M) and C57BL/6 mice (L).** Each row depicts a separate experiment, with the first columns depicting serum IgG against HA(CA07) (left) and HA(PR8) (right) (**A, D, G, J, K, L**). The next columns depict IgG1 and IgG2a titers against HA(CA07) and HA(PR8) before and after secondary exposure (**B, E, H),** and the last columns depict serum neutralization against influenza CA07 and PR8, respectively (**C, F, I**). **M**) Serum IgG against head domain HA1(CA07) and HA1(PR8) from experiment A.

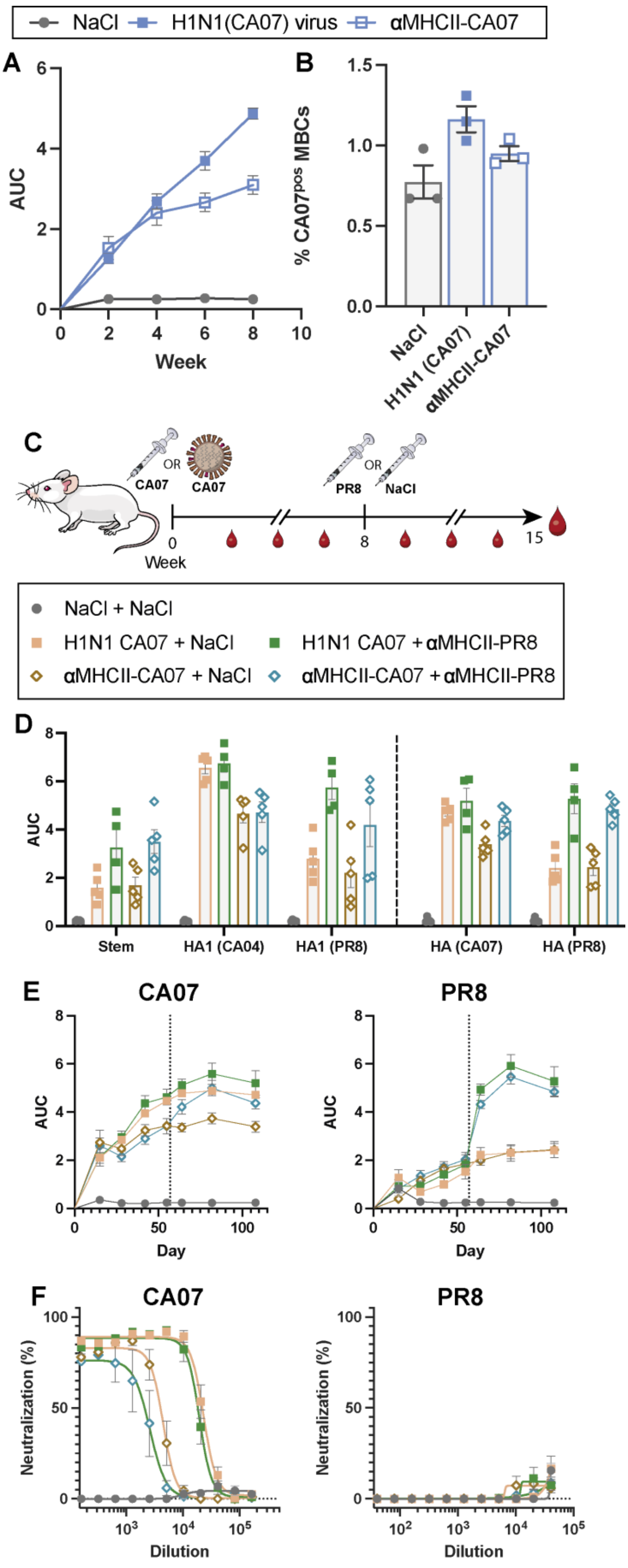
NaCl
H1N1(CA07) virus
αMHCII-CA07
A
AUC
Week
B
% CA07pos MBCs
NaCl
H1N1 (CA07)
αMHCII-CA07
C
CA07
OR
CA07
PR8
OR
NaCl
0
Week
8
15
NaCl + NaCl
H1N1 CA07 + NaCl
H1N1 CA07 + αMHCII-PR8
αMHCII-CA07 + NaCl
αMHCII-CA07 + αMHCII-PR8
D
AUC
Stem
HA1 (CA04)
HA1 (PR8)
HA (CA07)
HA (PR8)
E
CA07
PR8
AUC
Day
F
CA07
PR8
Neutralization (%)
Dilution

25

**Extended data figure 2: Sublethal viral influenza challenge or DNA vaccination will similarly induce antibody responses and imprinting in mice. A)** Mice were either vaccinated once with αMHCII-CA07 or infected with a sublethal dose of CA07 (n=5/group). Serum IgG antibodies against HA(CA07) are depicted. **B)** Mice were either vaccinated once with αMHCII-CA07, or infected with a sublethal dose of CA07 (n=3/group). Eighty days later, HA(CA07)$^{pos}$ MBC from spleen collected from the different groups were assessed and defined as: Viable single cells, TCRβ$^{neg}$ B220$^{pos}$ CD19$^{pos}$ CD45$^{pos}$ CD38$^{pos}$ PD-L2$^{pos}$. **C)** Experimental outline for D-F: Mice (n=5/group) were primed with either a sublethal dose (0.25×$LD_{50}$) of CA07 influenza virus or vaccinated with 50µg of αMHCII-CA07 DNA vaccine. On day 57 post priming, mice were given a secondary influenza exposure through vaccination with 50µg αMHCII-PR8 or mock (NaCl). **D)** IgG responses against the HA stem domain [shared between HA(CA07) and HA(PR8)], the HA1 domains of HA(PR8) and HA(CA07), as well as HA(CA07) and HA(PR8) were assessed in sera from day 108 after priming. **E)** IgG responses against HA(CA07) and HA(PR8) evaluated longitudinally after priming and secondary antigenic exposure. **F)** In vitro serum neutralization against H1N1 (CA07) (left) and H1N1 (PR8) (right) influenza viruses.

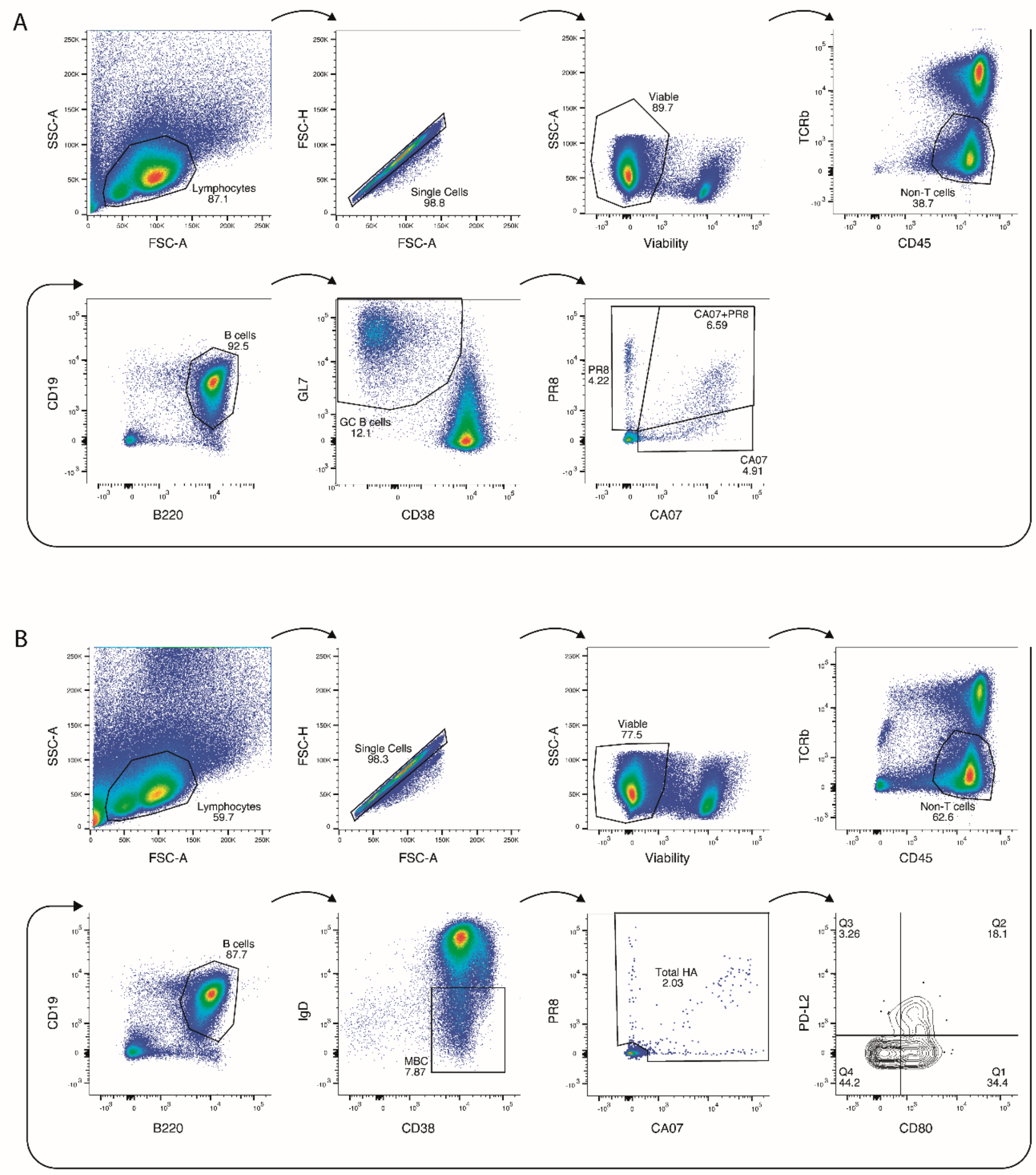


**Extended data figure 3**: **Gating strategies for GC B cells and MBC.** A) GC B cells were first gated as single and viable cells, and then as TCRb$^{neg}$ CD45$^{pos}$ CD19$^{pos}$ B220$^{pos}$ GL7$^{pos}$ CD38$^{neg}$. B) MBCs cells were first gated as single and viable cells, then TCRb$^{neg}$ CD45$^{pos}$

CD19$^{pos}$ B220$^{pos}$ IgD$^{neg}$ CD38$^{pos}$ HA$^{pos}$, and then either positive or negative for PD-L2 (CD273) and CD80.

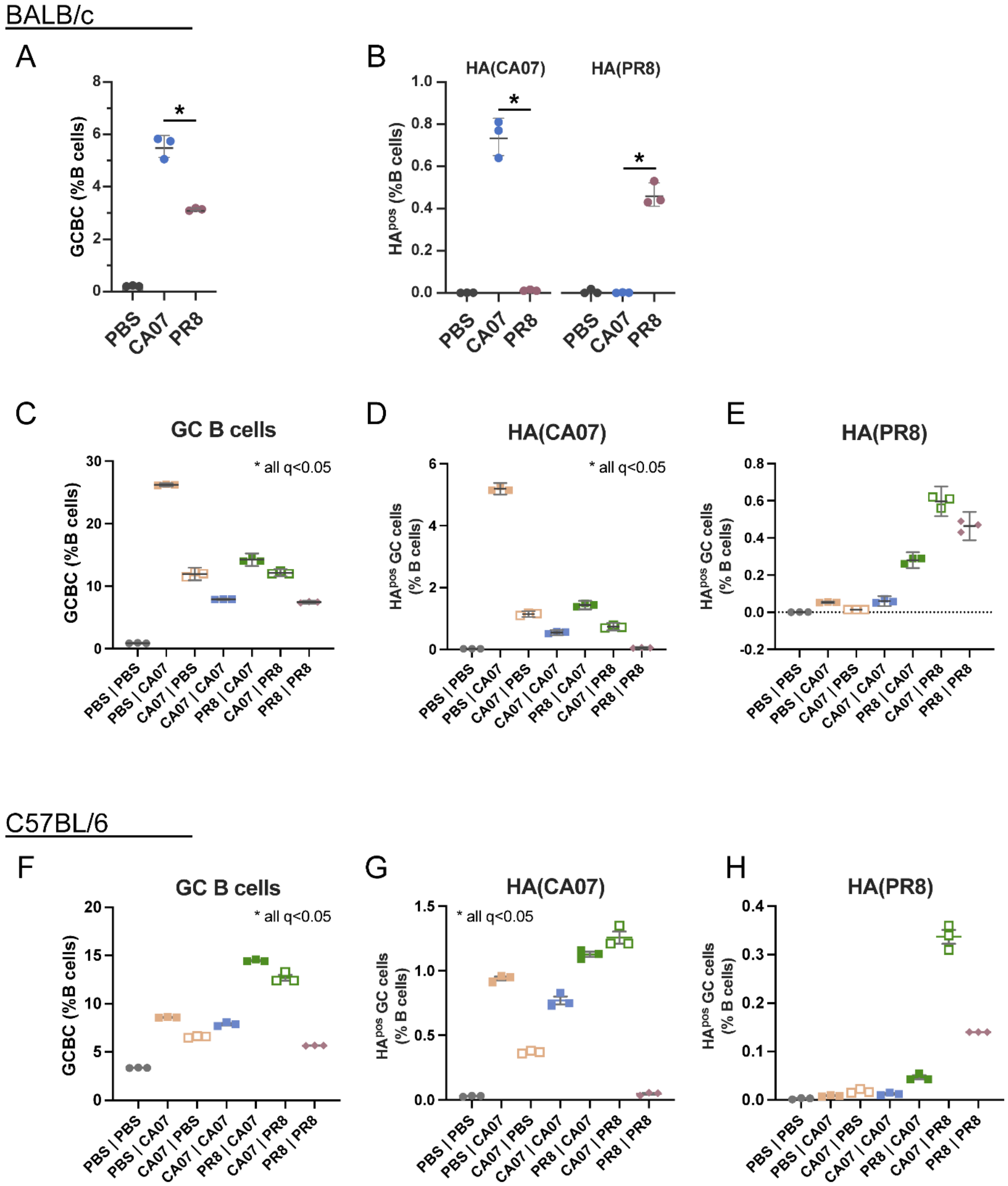


**Extended data figure 4: Data summaries of GC B cell evaluations.**

Depicted are summaries with quantifications corresponding to Fig. 2. In brief, mediastinal draining LN cells were collected and GC B cell evaluated by flow cytometry (gating strategy in Extended data Fig. 3A). **A-B)** GC B cells were collected from mice challenged sublethally (0.125×$LD_{50}$) with PR8, CA07, or NaCl, and defined as $GL7^{pos}$ and $CD38^{neg}$. **A)** Proportion of GC B cells depicted. **B)** Proportion of $HA^{pos}$ GC B cells depicted. **C-H)** GC B cells were collected from mice challenged twice with either homologous or heterologous H1N1 influenza viruses, as depicted. **C-E)** Summary of results from BALB/c mice challenged twice with sublethal doses if influenza virus (see Fig. 2C-D). **F-H)** Summary of results from C57BL/6 mice challenged twice with sublethal doses of influenza virus (see Fig. 2E-F). Statistical significance was calculated using one-way Brown-Forsythe and Welch ANOVA with false discovery rate using Benjamini, Krieger and Yekuteli method for multiple comparisons.

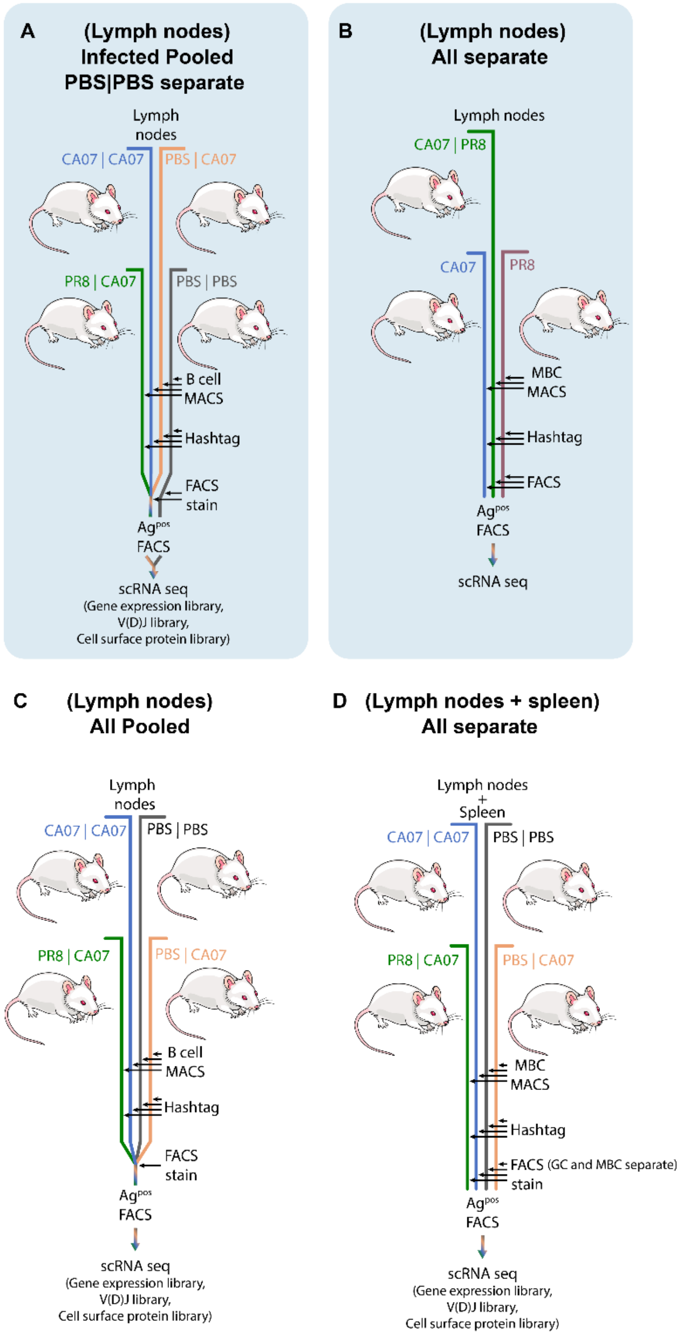


**Extended data figure 5**: **Illustration outlining organ harvest and FACS staining from 4 independent experiments with isolation of HA specific single B cells for BCR sequencing. A)** Draining LNs were harvested from infection groups (PBS+CA07, CA07+CA07, PR8+CA07), and B cells isolated for sequencing. B cells from uninfected mice (PBS+PBS) were sorted separately and combined with the remaining groups during GEM formation. **B)** Draining LNs were harvested from infection groups (CA07+PR8, CA07, PR8), and all infection groups sorted separately before combining during GEM formation. **C)** Draining LNs were harvested from infection groups (CA07+CA07, PR8+CA07, PBS+PBS, PBS+CA07). All groups were combined during surface staining but after hashtagging. **D)** Draining LNs and spleens were harvested from infection groups (CA07+CA07, PR8+CA07, PBS+PBS, PBS+CA07). Memory B cells were magnetically isolated from spleens before staining while no magnetic isolation was performed on draining LNs. All groups were sorted separately and combined during GEM formation.

Data from highlighted A and B are used for figures 4-5 and C and D are used in extended data figures 7-8.

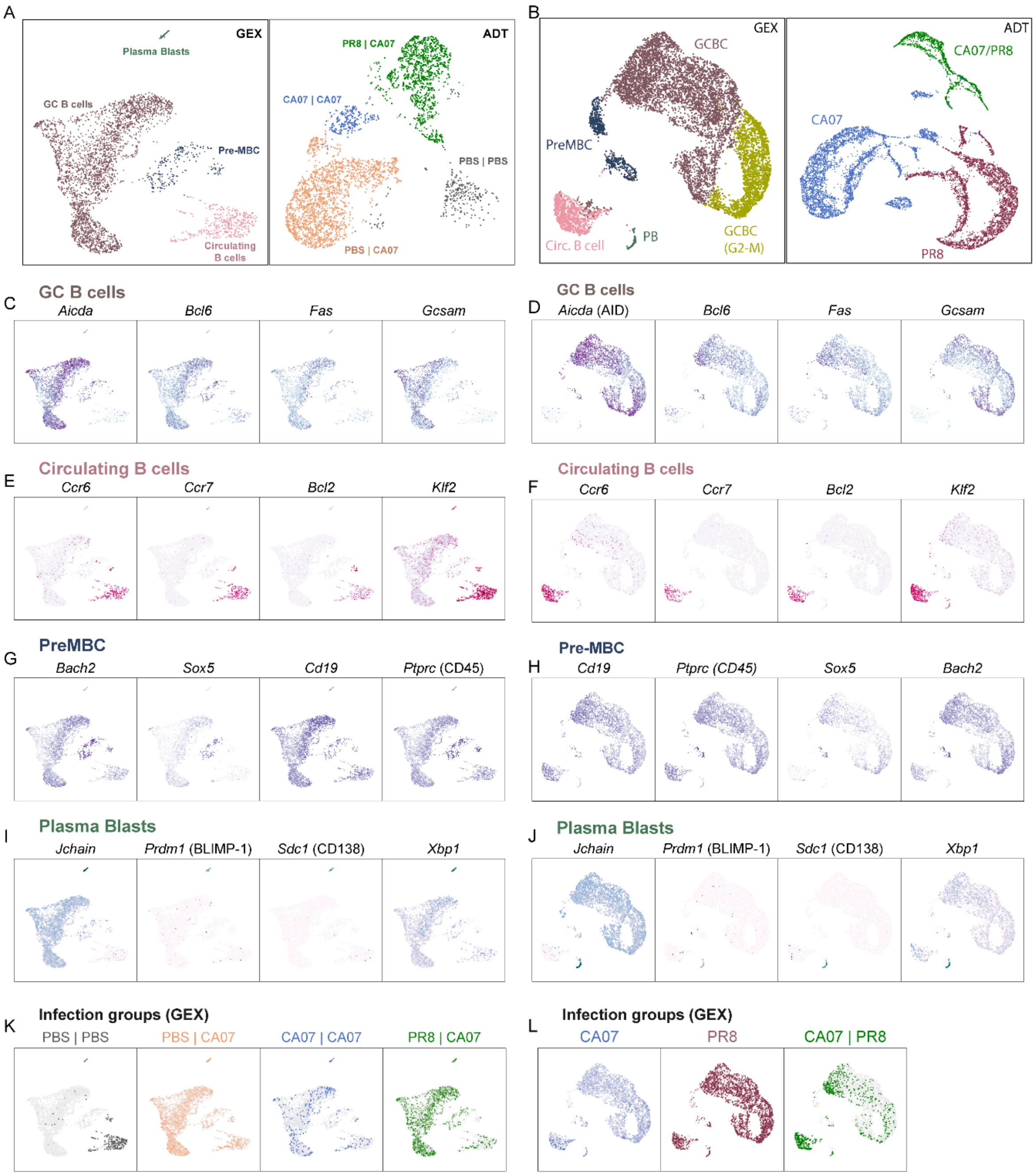


**Extended data figure 6: Additional GEX data from scRNA sequencing presented in Figure 4, GEX and ADT clustering. A)** GEX and **B)** ADT clustering of sequenced single HA-specific B cells. **C-D)** Example gene expression of identifying markers for GC B cells: *Aicda* (AID), *Bcl6*, *Fas* and Gcsam. **E-F)** Example gene expression of identifying markers for circulating B cells: *Ccr6*, *Ccr7*, *Bcl2*, *Klf2*. **G-H)** Example gene expression of identifying markers for pre-MBCs: *Bach2*, *Sox5*, *Cd19*, *Ptprc* (CD45). **I-J)** Example gene expression of identifying markers for plasmablasts: *Jchain*, *Prdm1* (BLIMP-1), *Sdc1* (CD138), *Xbp1*. **K-L**) The different infection groups were identified based on hashtag antibodies.

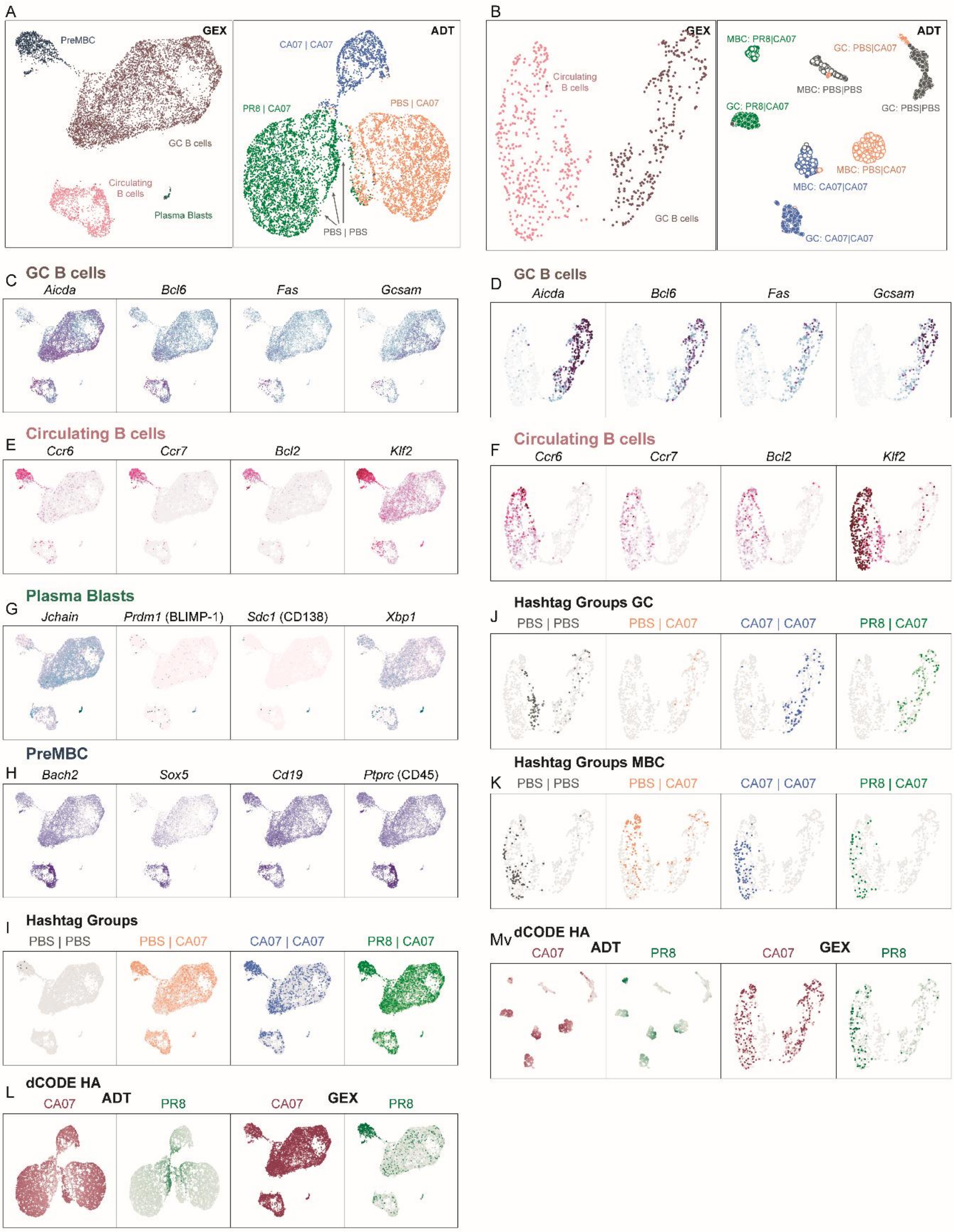


**Extended data figure 7: Additional GEX data from scRNA sequencing presented in Extended data figure 5C-D. A)** GEX and ADT clustering from Extended data Fig. 5C **B)** GEX and ADT clustering from Extended data Fig. 5D. **C-D)** Example gene expression of identifying markers for GC B cells: *Aicda* (AID), *Bcl6*, *Fas* and Gcsam. **E-F)** Example gene expression of identifying markers for circulating B cells: *Ccr6*, *Ccr7*, *Bcl2*, *Klf2*. **G)** Example gene expression of identifying markers for pre-MBCs: *Bach2*, *Sox5*, *Cd19*, *Ptprc* (CD45). **H)** Example gene expression of identifying markers for plasmablasts: *Jchain*, *Prdm1* (BLIMP-1), *Sdc1* (CD138), *Xbp1*. **I)** Infection groups were identified based on hashtag clusters for Extended data Fig. 5C. **J-K)** Infection groups were identified based on hashtag clusters for Extended data Fig. 5D. **L)** dCODE-HA binding based on reads per cell for Extended data Fig. 5C, displayed as both ADT and GEX cluster. **M)** dCODE-HA binding based on reads per cell for Extended data Fig. 5C, displayed as both ADT and GEX cluster.

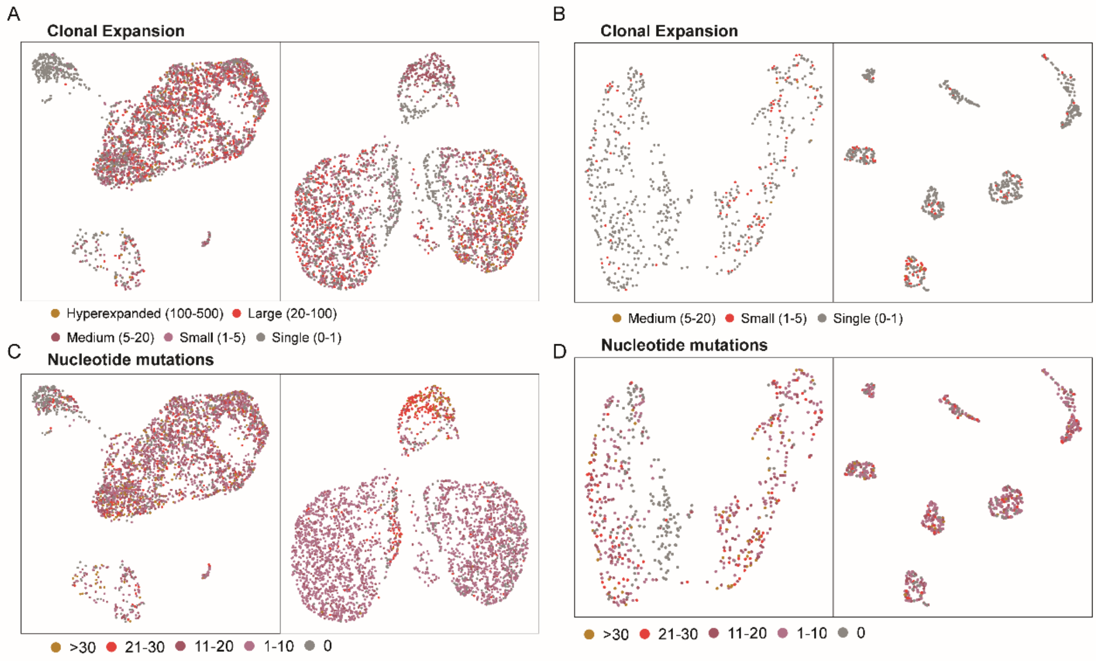


**Extended data figure 8: Clonal expansion and mutation analysis from Extended data figure 5C-D. A)** Clonal expansion analysis of Extended data Fig. 5C, shown on GEX and ADT clusters. **B)** Clonal expansion analysis of Extended data Fig. 5D, shown on GEX and ADT clusters. **C)** Mutational load analysis of Extended data Fig. 5C, shown on GEX and ADT clusters. **D)** Mutational load analysis of Extended data Fig. 5D, shown on GEX and ADT clusters.

| Clone | Exp | IGH GC | IGHV/IGLV | IGHD | IGHJ/IGLJ | CDRH3 | Count | HC CDR Mut. | Total Mut. |
|---|---|---|---|---|---|---|---|---|---|
| **SCA1** | E | IGHG 2A | BQCW - **IGHV1-69*02** | EVA3 – **IGHD2-8*01** | OMJ3 – **IGHJ3*01** | CARGRYGNTAFAYW | 34 | 0 | 2 |
| | | | 6QAU – **IGKV14-130*01 F** | | Z5J4 – **IGKJ4*01** | CLQFYEFPLTF | | | |
| **SCA2** | E | IGHG 3 | KPEP – **VBASE2: musIGHV021** | N42X – **IGHD2-2*01** | SCNX – **IGHJ4*01** | CARSPLVGGYYYPMDYW | 35 | 4 | 34 |
| | | | HA63 – **IGKV3-4*01 F** | | 4JXG – **IGKJ1*01** | CQQSNQDPWTF | | | |
| **SCA3** | D | IGHG 2A | O4YA - **IGHV3-2*02** | N.D. | ICAF – **IGHJ2*01** | CARSRGYFDYW | 74 | 0 | 0 |
| | | | 2UKG – **IGKV6-32*01 F** | | MXRV – **IGKJ2*01** | CQQDYSSPYTF | | | |
| **SCA4** | E | IGHG 2B | CFNH – **Haines: J558.6** | HNPT – **IGHD4-1*01** | G76U – **IGHJ1*01** | CARTGNWDGYFDVW | 1 | 25 | 72 |
| | | | GTW5 – **IGKV6-15*01 F** | | O4CL – **IGKJ5*01** | CQQYNSYPLTF | | | |
| **SPR1** | E | IGHG 2A | FMN7 – **IGHV3-6*02** | HNPT – **IGHD4-1*01** | SCNX – **IGHJ4*01** | CARGGLVYAMDYW | 7 | 2 | 14 |
| | | | KDB5 – **IGKV16-104*01 F** | | 4JXG – **IGKJ1*01** | CQQHNEYPWTF | | | |
| **SPR2** | D | IGHG 1 | UOVB – **IGHV14-3*02** | OWOX – **IGHD1-1*01** | OMJ3 – **IGHJ3*01** | CARSYGCCYEVAYW | 7 | 6 | 36 |
| | | | GTW5 – **IGKV6-15*01 F** | | 4JXG – **IGKJ1*01** | CQQYYSYPWTF | | | |
| **SPR3** | D | IGHG 1 | NNFU – **IGHV5-9-3*01** | EVA3 – **IGHD2-8*01** | ICAF – **IGHJ2*01** | CARPRGYGPFDYW | 4 | 10 | 25 |
| | | | G5I3 – **IGKV8-19*01 F** | | O4CL – **IGKJ5*01** | CQNDYAYPLTF | | | |
| **SPR5** | E | IGHG 2A | 5O75 – **JACKSON: balbIGHV010** | PF6D – **IGHD2-14*01** | ICAF – **IGHJ2*01** | CARRDRNYFDYW | 1 | 6 | 44 |
| | | | EVIF – **IGKV8-28*01 F** | | O4CL – **IGKJ5*01** | CQNDHHYPLTF | | | |
| **CR1** | E | IGHG 1 | CFNH – **Haines: J558.6** | N42X – **IGHD2-2*01** | SCNX – **IGHJ4*01** | CARVGPIYYGYDDYAMDYW | 2 | 8 | 32 |
| | | | UTLD – **IGKV4-57-1*01 F** | | O4CL – **IGKJ5*01** | CQQRSSYPLTF | | | |

| | | | | | | | | | |
|---|---|---|---|---|---|---|---|---|---|
| **SCA5** | D | IGHG2A | ML5T – **IGHV9-2-1*01** | OWOX – **IGHD1-1*01** | OMJ3 – **IGHJ3*01** | CARNYGNSPFAYW | 10 | 1 | 8 |
| | | | EN2O – **IGKV12-41*01 F** | | Z5J4 – **IGKJ4*01** | CQHFWSTPFTF | | | |
| **SPR4** | E | IGHG2A | 7BBU – **IGHV6-6*02** | EVA3 – **IGHD2-8*01** | OMJ3 – **IGHJ3*01** | CTGGGNYFAYW | 6 | 4 | 28 |
| | | | U5CY – **IGKV3-5*01 F** | | Z5J4 – **IGKJ4*01** | CQQSYEDPLTF | | | |

**Extended data Table 1**: Summary data and clonal information of the 11 monoclonal Abs that were initially chosen to be synthesized for *in vitro* validation. Including: Clone name, experiment (as shown in Extended data Fig. 1) the clone was isolated from, IgH isotype, V(D)J genes, CDR3 sequence, then number of identical clones in each clonotype, the total number of CDRH3 mutations, and the total number of mutations in both HC+LC, CDR+FR. The V(D)J gene names are provided as both "Label" based on Jackson et al. dataset from the OGRDB[71] and the corresponding IMGT or alternative name, also from the OGRDB. (N.D. = Not determinable)
